# BMAL2 controls adipose tissue inflammation and metabolic adaptation during obesity

**DOI:** 10.1101/2025.03.11.641984

**Authors:** Morgane A. Philippe, Blandine Fruchet, Lucie Cagninacci, Lucie Beaudoin, Alexis Gadault, Nicolas Venteclef, Etienne Challet, Agnès Lehuen, Ute C. Rogner, Amine Toubal

## Abstract

Contemporary lifestyle modifications such as changes in nutritional and sleep/wake rhythms increase the risk of metabolic and inflammatory complications linked to obesity, including type 2 diabetes (T2D) and metabolic dysfunction-associated steatohepatitis (MASH). BMAL2 (Brain and Muscle ARNT Like Protein 2) is a transcription factor belonging to the circadian clock transcriptional feedback loop which synchronizes internal biological rhythms to environment. In humans, reduced expression in white adipose tissue (WAT) and specific polymorphisms of *BMAL2* are associated with obesity and T2D. In this study we report that *Bmal2* invalidation in mice leads to increased body weight gain during diet-induced obesity. Loss of BMAL2 triggers the inflammatory response by increasing *Tnfα* expression and modifying adipocyte progenitor fate. This results in reduced lipid storage capacity within the WAT and increased ectopic storage in the liver. These functional and structural alterations culminate in the onset of hepatic steatosis and insulin resistance in liver and WAT. Overall, our investigations underscore the role of BMAL2 in the development and function of adipocytes, as well as in their inflammatory potential within the WAT. Our findings contribute to the understanding of the role of circadian clock genes in obesity and interconnected metabolic complications.

**Highlights:**

- The transcription factor BMAL2 is involved in metabolic complications of obesity in a mouse model of diet-induced obesity
- Invalidation of *Bmal2* worsens insulin resistance and hepatic steatosis induced by high fat diet
- Invalidation of *Bmal2* impairs visceral adipose tissue adaptation capacity in promoting inflammation and adipose progenitor decline

**Graphical abstract:** 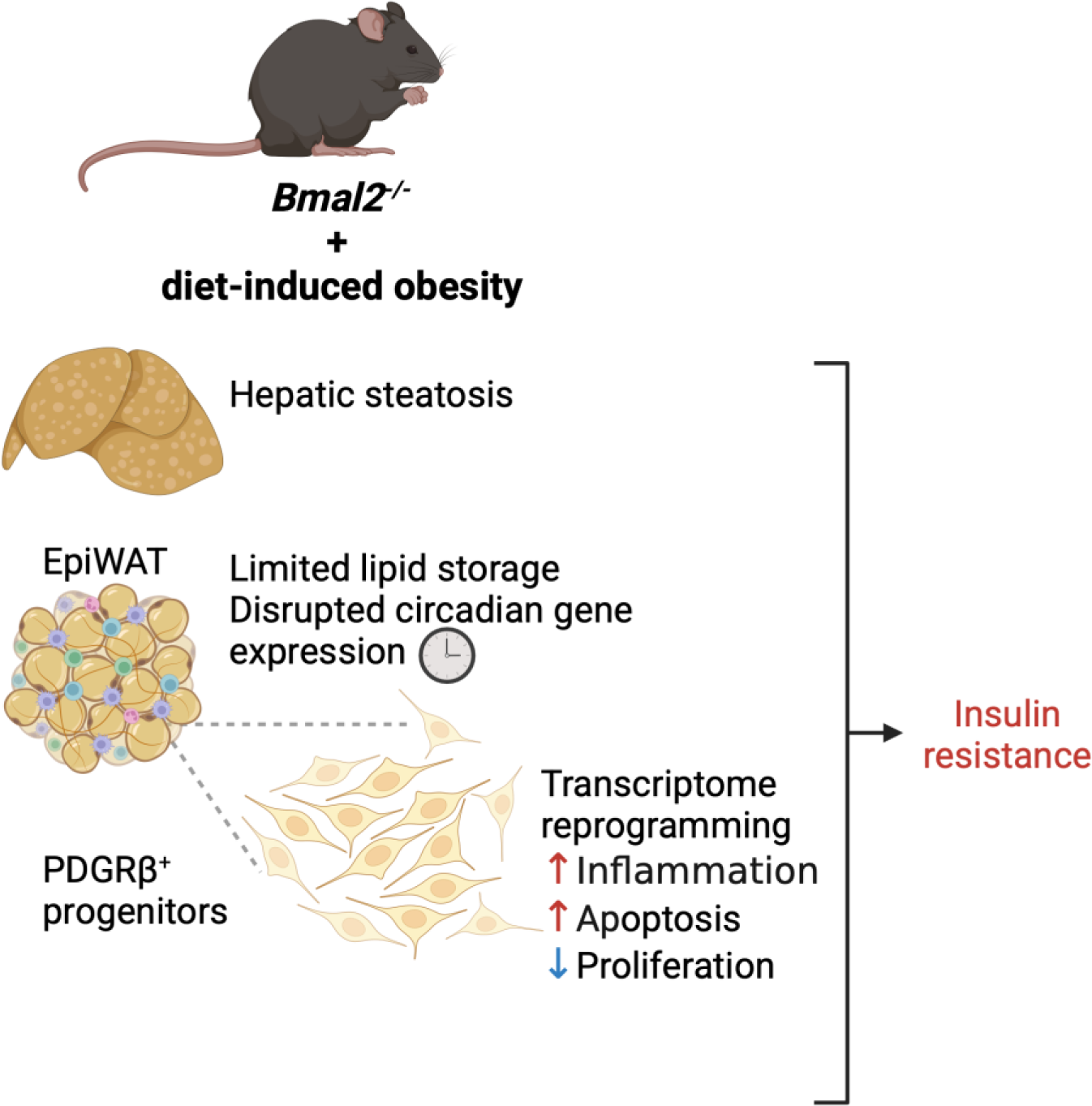

## 1. INTRODUCTION

The global emergence of obesity is now considered pandemic^1^. Primary drivers of this swift upsurge in obesity are the alterations of environment and lifestyle, notably manifested as disturbances in biological rhythms such as shifted work schedules, sleep patterns, and dietary habits^2^. In mammals, biological rhythms are controlled by the circadian clock, an intricate transcriptional feedback loop involving repressor (PER1/2:CRY1/2) and activator (BMAL1/2:CLOCK) elements^3,4^. The central clock, situated in the hypothalamus responds mainly to light, and controls downstream clocks in peripheral tissues that are sensitive to meal time^4,5^. Thus, rhythm disruption or mutations affecting circadian clock genes lead to modifications of behaviour, but also of internal metabolic functions^3,4,6^. Notably, desynchronization of the circadian system promotes obesity^3–6^. Circadian disruptions have for example been described in pancreatic islets^7^, peripheral immune cells^8^ and white adipose tissue^9,10^ of individuals afflicted by obesity and/or type 2 diabetes (T2D).

Obesity associated dysfunction of adipose tissue and chronic low-grade inflammation are causal to insulin resistance leading to the development of T2D^11,12^. The enduring low-grade inflammation arises from immuno-metabolic dysfunctions in mature adipocytes and adipose progenitors, alongside with the activation of tissue resident and infiltration of pro-inflammatory immune cells^13^. Adipocytes, with their ability to secrete adipokines governing metabolism and inflammation (such as leptin or adiponectin) in addition to chemokines, play a critical role in orchestrating the recruitment and activation of inflammatory cells into the tissue^14^. Under pathological conditions, adipocytes and their progenitors also produce inflammatory cytokines such as the tumor necrosis factor α (TNF-α) and interleukin 6 (IL-6) that inhibit adipocyte metabolic functions^15,16^ and have been directly linked to insulin resistance leading to T2D and hepatic steatosis^17,18^. Notably, TNF- α has first been identified as the major cytokine involved in adipose tissue low-grade inflammation^16,19^. Further studies highlighted the role of TNF-α in adipocyte dysfunctions^15,20,21^. Murine *Tnfα* knock-out models showed a pivotal role of TNF-α in the distribution of lipid deposition between adipose tissues and liver. *Tnfα* invalidation in mice enhances adipose tissue storage capacity, hepatic steatosis and therefore insulin sensitivity^22–26^.

As such, increased inflammation is associated with circadian dysfunction in mature adipocytes and adipose progenitors from obese patients^9^. While it has been established that circadian systems exert control over the storage capacity of adipose tissue^27^, the precise mechanisms governing circadian regulation of the inflammatory response in adipose tissue, particularly within adipocytes during obesity, remain poorly described. As a major clock transcription factor, BMAL1 (Brain and Muscle ARNT Like Protein 1, Basic Helix-Loop-Helix ARNT Like 1) related mechanisms are well known. BMAL1 is involved in glucose homeostasis control^28,29^ and metabolic adaptation during obesity and insulin resistance via the regulation of adipogenesis^30–32^, adipose tissue metabolic functions^33–36^ and compensatory pancreatic beta cells expansion^37,38^. *Bmal1* is an important regulator of inflammatory responses^39–41^, taking part in NF_k_B signalling pathway^9,42,43^ and in TNF-α production control in macrophages ^44–46^.

*Bmal2* (Basic Helix-Loop-Helix ARNT Like 2) is a paralog of *Bmal1*^47–49^. Like BMAL1, BMAL2 forms a transcriptionally active heterodimer with CLOCK and hypoxia-inducible factors^47^ and is involved in circadian transcriptional feedback loops^50^. Murine *Bmal2* is able to restore partly *Bmal1* functions in the control of obesity^51^. The role of *Bmal2* in physiology and disease may therefore be more important than has been appreciated to date.

In humans, several *BMAL2* polymorphisms have been associated with obesity and type 2 diabetes^52,53^, and *BMAL2* expression is downregulated in white adipose tissue of T2D patients^10^. *BMAL2* expression is regulated during weight-loss interventions in humans and has been shown to inhibit adipogenesis^54^. Our previous studies have delineated the role of *Bmal2* in inflammation during type 1 diabetes through the control of interleukin 21 expression in mice^55–59^. Hence, it is conceivable that BMAL2 might directly influence the regulation of inflammation associated with obesity. In recent mice studies, we have reported that a lack of *Bmal2* alters the circadian regulation of food consumption and energy expenditure^60^. Yet, the specific mechanisms by which impairment of BMAL2 expression or activity could lead to obesity and type 2 diabetes (T2D) are still to be thoroughly examined.

In the present study, using a C57BL/6 *Bmal2^-/-^* mouse model and the corresponding wildtype mice, we investigated metabolic and inflammatory responses in the liver and white adipose tissue during either a normal diet (ND) or high fat diet (HFD) feeding. We report that *Bmal2* deficiency impairs immune-metabolic functions in both mature adipocytes and their progenitors, leading to reduced lipid storage in epididymal white adipose tissue (EpiWAT). Concurrently, excess lipids accumulate ectopically in the liver, driving insulin resistance in both organs. Together, our findings underscore a key role of *Bmal2* in adipose tissue biology and highlight the broader importance of circadian regulation in obesity.

## 2. MATERIALS AND METHODS

### 2.1. Mice

The *Bmal2* mutant strain (designated by *Bmal2*^-/-^ or KO mice) carries the allele *Arntl2*^tm1a(KOMP)Wtsi^, has been imported from the UCDavis KOMP Repository (Davis, CA), and has been crossed to the C57BL/6J strain to obtain wildtype control (WT) and *Bmal2*^-/-^ strains^58^. Mice were fed a standard chow diet (ND, 12,6 kJ% fat, SAFE 150 SP-25). At 10 weeks of age, male mice were either fed with the standard chow diet or high fat diet (HFD, 60 kJ% fat, SSNIFF 813 ref# E15742-347) for 12 weeks. All mice were bred and maintained in the mouse facility of the Institut Pasteur, Paris under controlled temperature (21–23°C) and light (12-h light/12-h dark) conditions: lights were turned on at Zeitgeber time (ZT) 0 and turned off at ZT12. Depending on experiments, mice were sacrificed in the morning at ZT4 or at 6 time points over a complete 24-hour light-dark cycle (ZT0, ZT4, ZT8, ZT12, ZT16, ZT20).

### 2.2. Metabolic analysis

Oral glucose tolerance tests (OGTT) were performed after 10-12 weeks of diet as follows: overnight fasted mice were injected with glucose by gavage (2 g/kg body weight of glucose, 20% glucose solution; G8270, Sigma). Blood glucose concentration was determined at 0, 15, 30, 60, 90 and 120 min following the glucose load with a glucometer (Accu-Chek^®^ Performa, Roche). Blood samples from the tip of the tail vein were collected at 0, 15 and 30min following glucose load using Microvette CB 300 K2E (16.444, Sarstedt). Serum samples were collected through centrifugation to measure insulin level using Ultra-sensitive mouse insulin ELISA kit (90080, Crystal Chem).

Intraperitoneal (i.p.) insulin tolerance tests (ITT) were performed after 10-12 weeks of diet as follows: overnight fasted mice were injected with 0.00075 U of insulin/g of body weight (0.075 U/mL in PBS 1% BSA solution; Humalog 100 UI/mL, Lispro, Lilly). Blood glucose concentration was determined at 0, 15, 30, 60, 90 and 120 min with a glucometer (Accu-Chek^®^ Performa, Roche) on blood from the tip of the tail vein. The area under the curve (AUC, 0-120 min) was calculated for each group of mice.

For insulin-signaling assays, mice were fasted overnight and then i.p. injected with 1.00 U/kg insulin. Mice were euthanized 15 min after insulin injection, and liver and EpiWAT were collected for Western Blot analysis.

Lean tissue and fat mass of mice and liver biopsies were measured by nuclear magnetic resonance (NMR) using MiniSpec LF90 and miniSpec mq60, Bruker, according to the manufacturer’s instructions.

### 2.3. Histology

EpiWAT and liver samples were fixed in 4% formaldehyde solution overnight and embedded in paraffin. EpiWAT and liver slides were stained with hematoxylin/eosin or Sirius red for the evaluation of the tissue morphology, following standardized protocols. Adipocyte size was measured by the diameters of the adipocytes in light-microscopy images (×20) of EpiWAT sections (100-400 adipocytes per section, 1 section per mouse, 7-15 mice per group), and analyzed using ImageJ software.

### 2.4. Western Blot

Samples were lysed in RIPA buffer (Sigma) supplemented with protease (SIGMAFAST protease inhibitor tablets, Sigma) and phosphatase (PhosSTOP EASYpack, Roche) inhibitors, and were diluted with RIPA buffer, and Laemmli buffer (4X Laemmli sample buffer, Bio-Rad) and beta-mercaptoethanol to a concentration of 20 μg of protein and heated at 95 °C for 5 min. Proteins were separated by SDS–PAGE electrophoresis (4-12% Bis-Tris-Gel, NuPage Invitrogen) and transferred to 0.2µm nitrocellulose membranes (Bio-Rad). Blocking reagent (TBS with 0.5% Tween 20 and 3% BSA) were incubated for 1 h, and primary antibodies were incubated overnight at 4 °C in the blocking solution. The antibodies (listed in Supp.Table 1) and their concentrations were as follows: anti-phospho-AKT (P-AKT(S473) XP(R) Rabbit mAb, Cell Signaling 1:2000) and anti-beta-actin (Beta Actin Rabbit mAb, Cell Signaling 1:1000). After several washes in PBS with 0.5% Tween 20, horseradish peroxidase (HRP)-labeled secondary antibodies (Anti-rabbit IgG HRP-linked Antibody, Cell Signaling 1:2000) were incubated for 1 h at room temperature in the blocking solution. Membranes were incubated with ECL western-blotting substrate (ECL Prime Western Blotting Detection Reagents, Amersham, Cytiva) and imaged by the myECL Imager (ThermoFisher) or FUSION FX (Vilber). Blots were semi-quantified using ImageJ software.

### 2.5. Immune cell isolation and flow cytometry

The EpiWAT fat pad was isolated from mice and digested with 5 ml of collagenase H (2 mg/ml, Roche) and BSA (2.5%, Sigma) solution at 37 °C for 30 min with shaking (150 rpm). After digestion, adipocytes were removed by filtering through a 100 μm cell strainer, and cell suspension was centrifuged for 5 min at 300 g to pellet the stroma-vascular fraction (SVF). Red blood cells in the SVF pellet were lysed by brief incubation in 1 ml RBC lysis buffer (Sigma). SVF was washed with PBS containing 5% fetal calf serum (FCS) and 0.1% sodium azide in PBS. Macrophages were directly analyzed from SVF, whereas other immune cells were further enriched on Percoll density gradients of 40 and 80% (GE Healthcare). The interface between the layers was collected and suspended in PBS containing 5% FCS and 0.1% sodium azide, to retrieve immune cells.

Liver was perfused with RPMI 1640 medium (Gibco) supplemented with 5% FCS to remove circulating blood cells and then harvested. Liver was passed through 70 μm cell strainer. Cells suspension was collected after 2 min centrifugation at 48 *g* to avoid parenchymal cells and was centrifuge again at 440 *g*. Red blood cells were lysed by brief incubation in 1 ml RBC lysis buffer (Sigma), centrifuge again at 440 *g* and the pellet was then resuspended in 40% Percoll layered onto 80% Percoll and centrifuged for 15 min at 780 *g* at room temperature. The immune cell fraction was collected at the interface and suspended in PBS containing 5% FCS and 0.1% sodium azide, to retrieve immune cells.

Cell suspensions prepared from various tissues were labeled at 4 °C in PBS containing 5% FCS and 0.1% sodium azide. Surface labeling was performed at 4°C during 30 min with the following antibodies: CD45 APC-Cy7, Ly6C BV510, CD206 FITC, F4/80 PE-Cy7, CD11c APC, TCRb BV711 BD, CD11b BV785, TCRgd Percp-Cy5.5, CD4 BV510, CD8 BV605 (listed in Supp.Table 1).

Intra-cytoplasmic labeling for cytokine analysis were performed after PMA (25 ng/ml) and ionomycin (1 μg/ml) stimulation for 4 h at 37 °C in presence of Brefeldin A (10 μg/ml), all reagents from Sigma-Aldrich. After surface labeling, cells were fixed and permeabilized with Cytofix/Cytoperm kit (BD Biosciences), washed, and incubated at 4 °C with antibodies: TNF-α BV421, IFNγ AF700, IL17-A Percp-Cy5.5 (listed in Supp.Table 1). Data acquisition was performed using a BD Biosciences LSR Fortessa cytometer then the results were analyzed using FlowJo analysis software (Tree Star).

### 2.6. Adipose progenitor isolation and flow cytometry

Adipose progenitor preparation and isolation were adapted from Hepler et al. 2018^61^. The EpiWAT fat pad was isolated from mice and digested with 5 ml of collagenase H (2 mg/ml, Roche) and BSA (2.5%, Sigma) solution at 37 °C for 45-60 min with shaking (180 rpm). The solution of digested tissue was passed through a 100 µm cell strainer, diluted to 30 ml with RPMI 1640 medium (Gibco) supplemented with 5% FCS, and centrifuged at 300 *g* for 5 min. The supernatant was aspirated and red blood cells in the SVF pellet were lysed by brief incubation in 1 ml RBC lysis buffer (Sigma). Next, the mixture was diluted to 10 ml with RPMI 1640 medium (Gibco) supplemented with 5% FCS, passed through a 40 μm cell strainer, and then centrifuged at 500 *g* for 5 min. The supernatant was aspirated, and cells were resuspended in PBS containing 5% FCS, 0.1% sodium azide and 1% BSA. Cell suspensions were stained at 4 °C in PBS containing 5% FCS, 0.1% sodium azide and 1% BSA. Surface staining was performed at 4°C during 20 min with the following antibodies (listed in Supp.Table 1): CD31 Percp-Cy5.5 1/400, PDGFRβ PE 1/50, CD9 APC 1/400, CD45 APC-Cy7 1/200, Ly6C BV510 1/100.

Data acquisition was performed using a BD Biosciences LSR Fortessa cytometer, and cell sorting was performed using BD Biosciences FACSAria III, then the results were analyzed using FlowJo analysis software (Tree Star).

### 2.7. Adipose progenitor cell isolation, siRNA transfection and 3D spheroid differentiation

Adipose progenitor cells were isolated from the EpiWAT of C57BL/6 mice. Briefly, the tissue was digested in collagenase H, and the resulting SVF was collected by centrifugation. The SVF was then seeded in DMEM (Thermo Fisher #10566016) supplemented with 10% fetal bovine serum (FBS) and penicillin/streptomycin (PS), and cultured for 7 days to allow cell expansion.

Following this expansion phase, cells were harvested and counted. For each well, 50,000 cells were resuspended in antibiotic-free DMEM and transfected with 50 nM si*Bmal2 (*5’-AAGCTGGCCTCTGAATGTTGT-3’, Eurofins) or non-targeting control siRNA (siCtrl, 5’-AGGUAGUGUAAUCGCCUUG-3’, Eurofins) using Lipofectamine™ 3000 (Thermo Fisher #L3000001) at a 1:3 reagent-to-siRNA ratio, according to the manufacturer’s protocol. After 2 hours of incubation with the transfection complexes, the medium was replaced with fresh DMEM (10% FBS, 1% PS). Subsequently, the transfected cells were transferred to 96-well low-attachment plates (In-Sphero #CS-PB15) at 50,000 cells per well. Cells in control condition, without treatment, were directly seeded without transfection. Cells were then cultured for 48 hours in a seeding medium composed of DMEM, 10% FBS, 1% PS, and 0.1% fungizone. To initiate adipocyte differentiation, the medium was replaced with an adipogenic induction medium (DMEM, 10% FBS, 1% PS, 0.1% fungizone, 5 μg/mL insulin, 50 μM isobutylmethylxan-thine, 1 μM dexamethasone, 1 μM rosiglitazone) and maintained for 72 hours. Following this induction period, the cells were switched to a maintenance medium (DMEM, 10% FBS, 1% PS, 0.1% fungizone, 5 μg/mL insulin) for an additional 5 days.

In total, the cells underwent 8 days of differentiation, resulting in the formation of 3D spheroids, which were then harvested for further analyses. Knockdown efficiency was verified by quantitative RT-PCR (qRT-PCR) for target gene expression 48 hours post-transfection, confirming a robust reduction in *Bmal2* mRNA levels. Non-targeting control siRNA-treated cells were used to account for non-specific effects of the transfection reagent.

### 2.8. RT-qPCR

RNA was extracted from tissues or cultured cells using the RNeasy RNA Mini Kit (RNeasy mini Kit, Qiagen). RNA from cell sorted adipose progenitors was extracted with RNeasy RNA Plus micro Kit (RNeasy Plus micro Kit, Qiagen). Complementary DNAs were synthesized using Superscript III reverse-transcriptase kit (Invitrogen). Quantitative PCR analysis was performed with SYBR Green Master Mix (Roche) using a LightCycler 480 (Roche). *18S* was used for normalization to quantify relative mRNA expression levels. Relative changes in mRNA expression were calculated using the comparative cycle method (2^−ΔΔCt^). Primers are listed in Supp.Table 2.

### 2.9. Bulk RNA-seq: preparation, sequencing and analysis

Adipose progenitors defined as CD45^-^CD31^-^PDGFRβ^+^ were flow sorted and frozen following RNeasy RNA Plus micro Kit (Qiagen) instructions to extract RNA. Quality of the RNA was measured using a Bioanalyzer 2100 (Agilent) as per the manufacturer’s instructions using pico chip kit, RIN > 6,5 were kept for sequencing.

Non-strand-oriented libraries were prepared following the NEBNext Single Cell/Low Input RNA Library Prep Kit (New England BioLabs), paired end (2 x75 bp) sequencing was performed on an Illumina Nextseq 500 platform.

Fastq files were then aligned using STAR algorithm (version 2.7.6a), on the Ensembl release 101 GRCm38 reference. Reads were then count using RSEM (v1.3.1) and the statistical analyses on the read counts were performed with R (version 3.6.3) and the DESeq2 package (DESeq2_1.26.0) to determine the proportion of differentially expressed genes between two conditions. We used the standard DESeq2 normalization method (DESeq2’s median of ratios with the DESeq function), with a pre-filter of reads and genes (reads uniquely mapped on the genome, or up to 10 different loci with a count adjustment, and genes with at least 10 reads in at least 3 different samples).

Following the package recommendations, we used the Wald test with the contrast function and the Benjamini-Hochberg FDR control procedure to identify the differentially expressed genes. R scripts and parameters are available on the platform https://github.com/GENOM-IC-Cochin/RNA-Seq_analysis. Differentially expressed genes were defined with log2FC>0.6 and p-adj<0.05.

### 2.10. GSEA Analysis

Gene set enrichment analysis (GSEA, Broad Institute) was done on hallmark genes. Database used: Mh.all.v2023.1 using mouse symbols; 1000 permutations, no collapse, permutation type was phenotype; max size of data set 500, min size 15.

### 2.11. Statistics

Statistical analyses were performed with GraphPad Prism 8.3.0 (GraphPad Software, Inc., La Jolla, CA) or R software 4.1.0. by two-tailed Mann-Whitney test and Spearman correlation test. To analyze and compare circadian gene expression rhythms in mice tissues, we used Circacompare package using R software 4.1.0. to assess rhythmicity for each data set and statistical differences between cosinor parameters (Phase, MESOR, Amplitude) of *Bmal2*^-/-^ and control mice. All data were represented as mean ± S.E.M, *P* < 0.05 was defined as significant.

### 2.12. Study approval

All animal studies have been approved by the relevant institutional review boards (Comité d’Ethique en Expérimentation Animale CETEA, IDF, Paris and C2EA 89, Paris) and the French ministry of higher education and research (Ministère de l’enseignement supérieur et de la recherche) under the reference number APAFIS#31813-202105271511696 v1.

## 3. RESULTS

### 3.1. *Bmal2* invalidation enhances body weight gain and reshapes lipid storage distribution during diet-induced obesity

We subjected ten-week-old male C57BL/6 *Bmal2*^-/-^ (KO) mice and age-matched wild type controls (WT) to either ND or HFD.

In order to first explore whether obesity impact *Bmal2* expression in WT controls, we measured its expression over 24-hour-light-dark cycle in liver and EpiWAT from both ND and HFD WT fed mice. In HFD condition, *Bmal2* displayed a significantly altered expression pattern across the 24-hour cycle compared to ND. Specifically, the midline estimation statistics of the rhythm (MESOR) increased significantly in the liver while in EpiWAT, *Bmal2* gained a marked 24-hour expression rhythm. This was accompanied by significant increase in both the MESOR and amplitude of its rhythmic profile (Figure 1A).

**Fig. 1.**
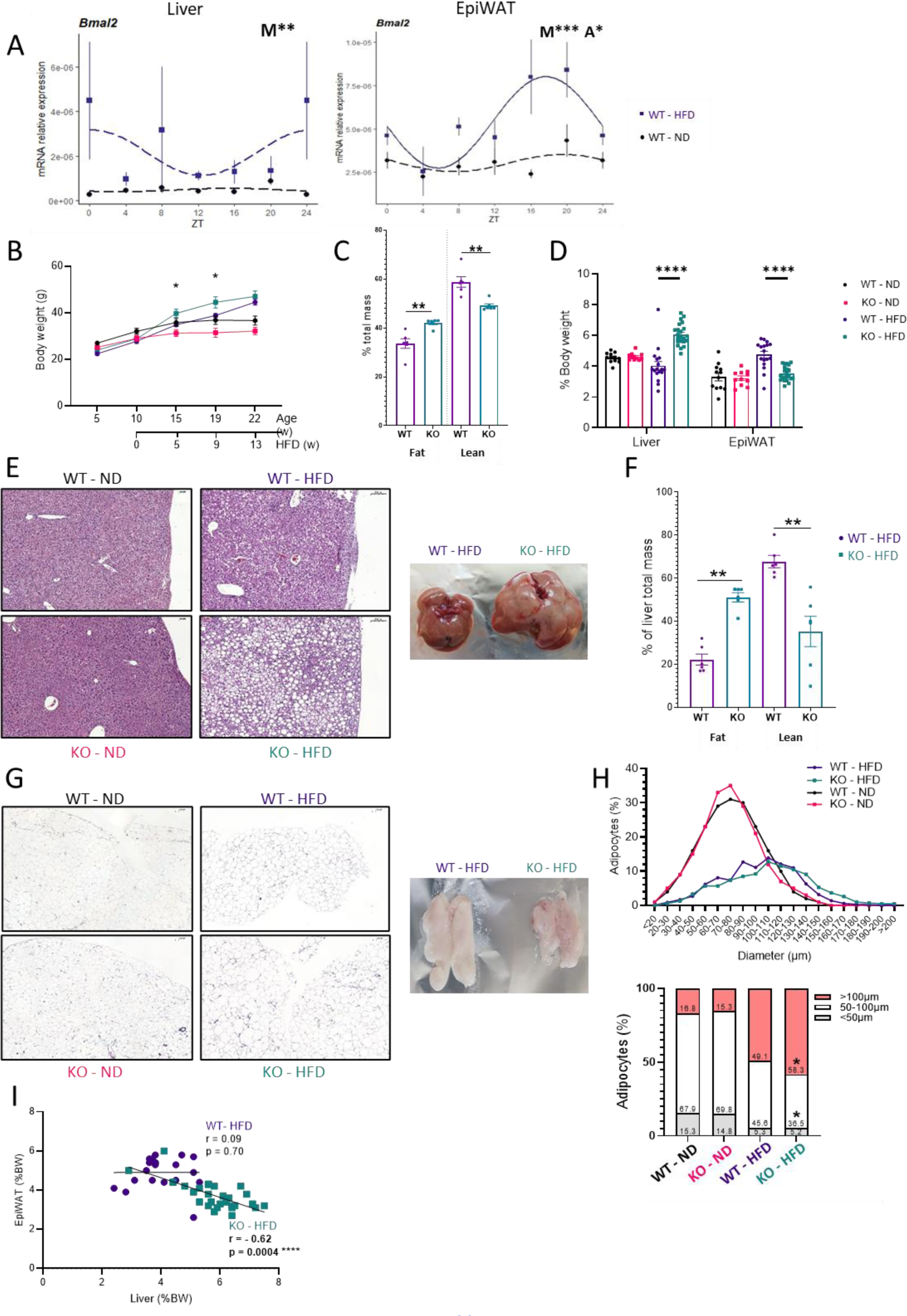
*Bmal2* invalidation induces increased body weight gain and modified lipid storage distribution during diet-induced obesity. A) Expression of *Bmal2* across a 24h cycle in liver and EpiWAT of control mice (WT) fed by ND or 12 weeks of HFD (n=3-8 per group). Circacompare package was used to assess rhythmicity for each data set and statistical differences between cosinor parameters (M: Mesor; A: Amplitude; *P < 0.05, **P < 0.01, ***P < 0.001). Data are represented as mean ± S.E.M with a line representing the estimated cosinor model. Solid line = significant rhythmicity, dashed line = no rhythmicity. B) Body weight (KO-ND n=3-5, WT-ND n=6, KO-HFD n=6, WT-HFD n=7), C) NMR-evaluated body fat and lean mass (KO-HFD n=6, WT-HFD n=6), D) Weight of liver and EpiWAT (KO-ND n=11, WT-ND n=12, KO-HFD n=22, WT-HFD n=17) relative to body weight (%), E) NMR-evaluated liver fat mass (KO-HFD n=6, WT-HFD n=6), E) Representative haematoxylin/eosin staining (scale bars, 100 μm) and pictures of liver and F) NMR-evaluated liver fat and lean mass (KO-HFD n=6, WT-HFD n=6), G) Representative haematoxylin & eosin staining of EpiWAT (scale bars, 200 μm) and representative pictures of EpiWAT, H) Quantification of adipocyte size in EpiWAT (100-400 adipocytes per section, 1 section per mouse, 7-15 mice per group), and I) Spearman correlation between weight of EpiWAT and liver relative to body weight (%) from *Bmal2^-/-^* mice (KO) and controls (WT) fed with ND and/or HFD during 12 weeks, as detailed on the figure. All data are represented as mean ± S.E.M. All statistical analyses were performed by two-tailed Mann-Whitney test. *P < 0.05, **P < 0.01, ***P < 0.001.

Then, we evaluated the phenotype of *Bmal2*^-/-^ mice to investigate the role of *Bmal2* in obesity and evaluate its influence on the disease. *Bmal2*^-/-^ mice exhibited greater weight gain than WT mice when fed HFD (Figure 1B and Supp. Figure 1A). NMR analyses further indicated an increase in total body fat mass in *Bmal2*^-/-^ mice compared to WT mice (Figure 1C). Liver weight measurements under HFD condition were significantly higher in *Bmal2*^-/-^, whereas EpiWAT weight was lower relative to WT (Figure 1D). In contrast, subcutaneous white adipose tissue (ScWAT) weight was significantly increased in *Bmal2*^-/-^ mice compared to WT mice under HFD (Supp.Figure 1B). Notably, no differences in organ weights were detected between *Bmal2*^-/-^ and WT mice fed with ND (Figure 1D and Supp.Figure 1B).

To probe the structural alterations induced by *Bmal2* deficiency, we next examined the liver and EpiWAT. Histological and NMR analyses of the liver revealed marked ectopic lipid accumulation in *Bmal2*^-/-^ mice under HFD conditions (Figure 1E-H). In contrast, there was no significant increase in muscle fat mass (Supp.Figure 2E). Adipocyte diameters in EpiWAT were measured using histological sections (Figure 1G-H). Under HFD, *Bmal2*^-/-^ mice showed a significantly higher percentage of large adipocytes (>100µm diameter) and a lower percentage of intermediate-sized adipocytes (50-100µm diameter) than WT mice, indicating EpiWAT adipocyte hypertrophy in *Bmal2*^-/-^ mice (Figure 1H). Both adipocyte diameter and fat pad mass were elevated in ScWAT of *Bmal2*^-/-^ mice on HFD, suggesting both hypertrophic and hyperplastic growth in this depot (Supp.Figure 1B-D).

**Fig. 2.**
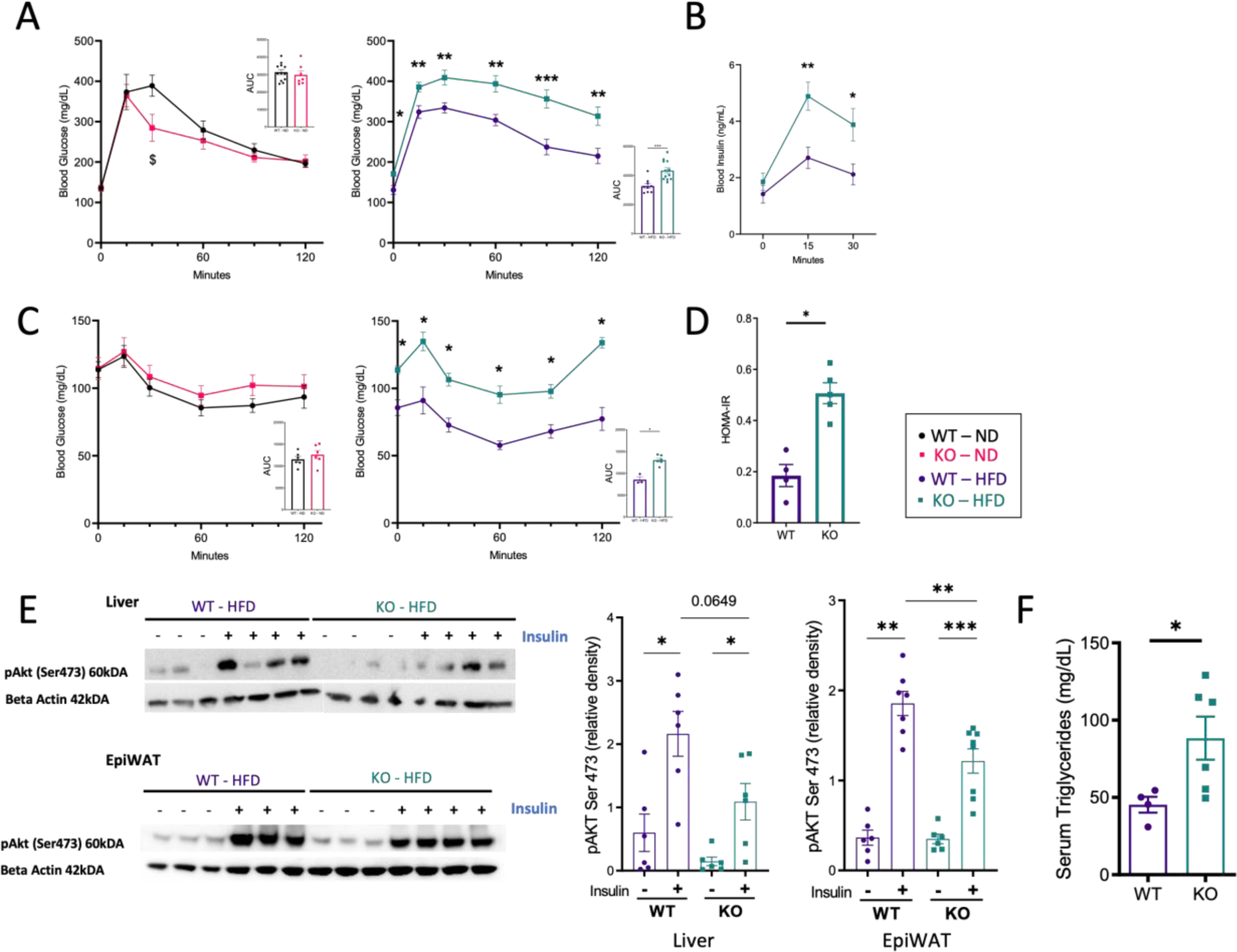
*Bmal2* invalidation enhances insulin resistance during diet-induced obesity. A) OGTT (KO-ND n=7, WT-ND n=5, KO-HFD n=12, WT-HFD n=9). Pooled data from two independent experiments are represented, B) Plasma insulin during OGTT (n=9-12 per group). Pooled data from two independent experiments are represented, C) ITT (KO-ND n=6, WT-ND n=6, KO-HFD n=6, WT-HFD n=3), D) HOMA-IR index (WT-HFD n=4, KO-HFD n=5), E) Western blot (left) and quantification (right) of AKT phosphorylation (p-AKT-S473) and β-actin in liver and EpiWAT after intraperitoneal insulin administration (1 UI/kg) (quantification n=6-8 mice per group), and F) Serum triglycerides (mg/dL) (WT-HFD n=4, KO-HFD n=6) in *Bmal2^-/-^* mice (KO) and controls (WT) fed with ND and/or HFD during 12 weeks. All data are represented as mean ± S.E.M. All statistical analyses were performed by two-tailed Mann-Whitney test. *P < 0.05, **P < 0.01, ***P < 0.001.

A correlation analysis between liver weight and EpiWAT weight revealed a negative association in *Bmal2*^-/-^ mice fed HFD, but not in WT mice (Figure 1I). This finding suggests a direct role of *Bmal2* in regulating lipid storage capacity within adipose tissues and in preventing ectopic lipid deposition in the liver under obesogenic conditions. Importantly, no such correlations were observed under ND (Supp.Figure 1F-G).

### 3.2. *Bmal2* invalidation promotes insulin resistance in the context of obesity

Next, we conducted oral glucose tolerance tests (OGTT) and insulin tolerance tests (ITT) in *Bmal2*^-/-^ and WT mice fed ND and HFD. Under HFD conditions, *Bmal2*^-/-^ mice exhibited significantly impaired glucose tolerance was observed in compared to WT, whereas no differences were detected under ND (Figure 2A). The observed glucose intolerance in *Bmal2*^-/-^ mice was accompanied by elevated insulin secretion (Figure 2B) indicating decreased insulin sensitivity in *Bmal2*^-/-^ mice under HFD. In agreement, obese *Bmal2*^-/-^ mice showed higher blood glucose levels during ITT than WT mice (Figure 2C), and a concomitant increase in the homeostasis model assessment of insulin resistance (HOMA-IR) index (Figure 2D).

To further assess insulin signaling, we measured Ser473-Akt phosphorylation (pAKT-Ser473), a key read-out for intracellular insulin signaling. Under HFD, *Bmal2*^-/-^ mice exhibited significantly reduced pAKT-Ser473 levels in both liver and EpiWAT compared to WT (Figure 2E). Consistent with these findings, obese *Bmal2*^-/-^ mice displayed elevated circulating triglyceride levels (Figure 2F), potentially reflecting impaired insulin-mediated suppression of lipolysis.

To determine whether *Bmal2* deficiency further affected EpiWAT function, we measured the expression of genes involved in adipogenesis (*Pparg*), adipose tissue homeostasis function (*Leptin*, *Adipoq*), and lipolysis (*Hsl*, *Atgl*). While no statistically differences were observed (Supp.Figure 2A), *Pparg* and *Adipoq* were slightly decreased and *Leptin* slightly increased in the EpiWAT of *Bmal2*^-/-^ mice on HFD. Similarly, the expression of key hepatic fatty acid synthesis genes (*Acly*, *Acaca*, *Scd1*, *Elovl6*) was not markedly altered in liver by *Bmal2* deficiency (Supp.Figure 2B). Taken together these findings suggest that despite limited changes in metabolic gene expression, *Bmal2* invalidation in obesogenic conditions leads to increased insulin resistance and altered lipid handling.

### 3.3. *Bmal2* invalidation exacerbates inflammation in the liver and visceral adipose tissue during diet-induced obesity

We next investigated the expression of key genes associated with metabolic inflammation in *Bmal2*^-/-^ and WT mice. In liver of *Bmal2*^-/-^ ND-fed mice, *Tnfα* expression was increased compared to WT mice (Figure 3A). The difference became even more pronounced with HFD feeding reaching 6-fold increase in obese *Bmal2*^-/-^ mice compared to obese WT mice.

**Fig. 3.**
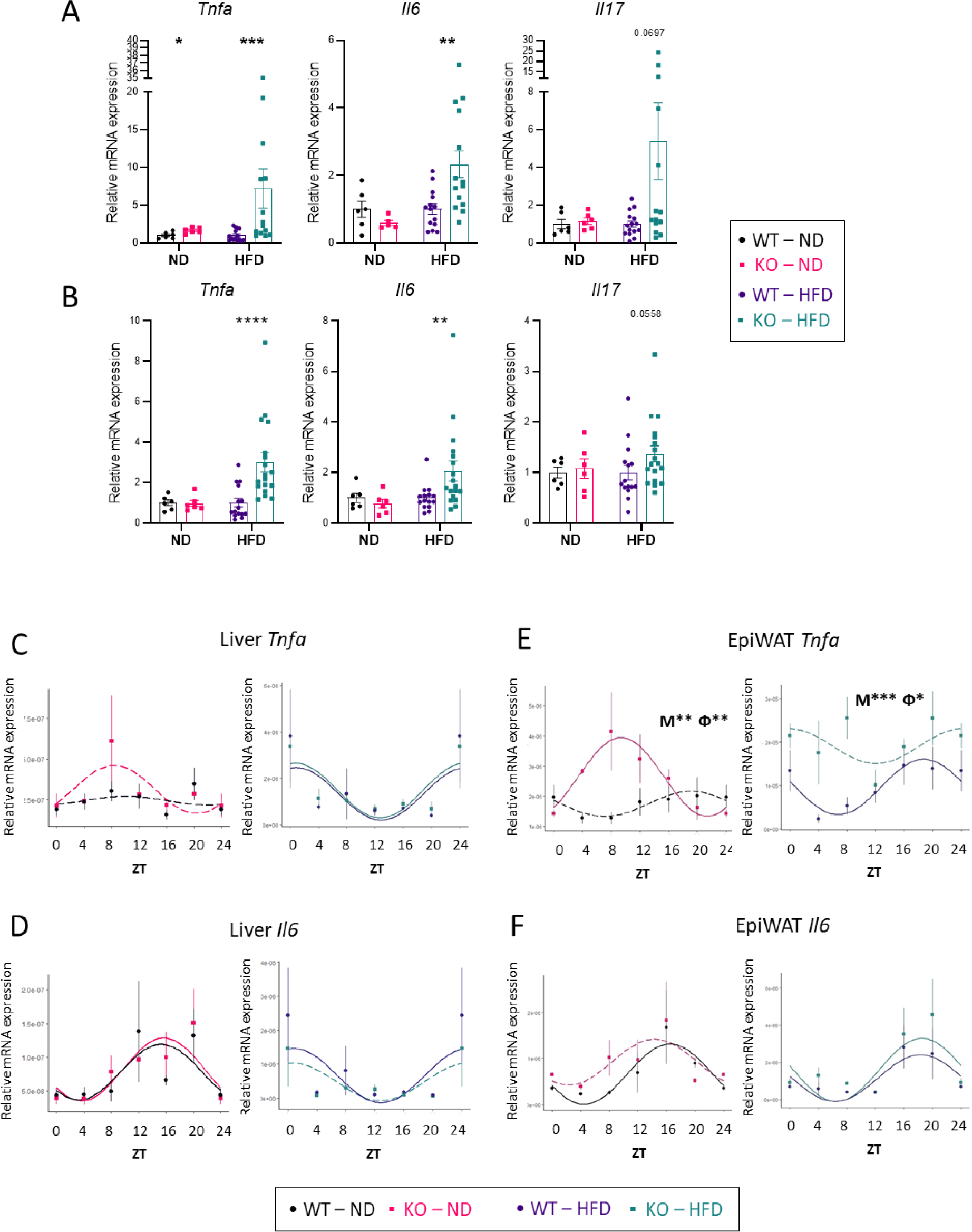
Increased inflammatory gene expression in liver and EpiWAT of obese *Bmal2^-/-^* mice. A-B) RT-qPCR analysis of inflammatory (*Tnfa*, *Il6*, *Il17*) gene expression in (A) liver (n=5-16 per group) and (B) EpiWAT (n=6-18 per group) from *Bmal2^-/-^*mice (KO) and controls (WT) fed with ND or HFD during 12 weeks). Data are represented as mean ± S.E.M. Statistical analyses were performed by two-tailed Mann–Whitney test. *P < 0.05, **P < 0.01, ***P < 0.001. B) RT-qPCR analysis of inflammatory gene expression (*Tnfa*, *Il6*, *Il17*) and C-F) of *Tnfa*, *Il6* across a 24h cycle in liver (C,D) EpiWAT (E,F) from *Bmal2^-/-^* mice (KO) and controls (WT) fed with ND or HFD during 12 weeks (n=3-8 per group). Circacompare package was used to assess rhythmicity for each data set and statistical differences between cosinor parameters (M: Mesor; A: Amplitude; Ф: Phase, *P < 0.05, **P < 0.01, ***P < 0.001). Data are represented as mean ± S.E.M with lines representing the estimated cosinor model. Solid line = significant rhythmicity, dashed line = no rhythmicity.

Similarly, *Il6* expression was significantly elevated in the liver of *Bmal2*^-/-^ mice under HFD, while *Il17* was only slightly increased compared to WT. Under ND, no significant differences were observed in *Il6* and *Il17* expression (Figure 3A). In EpiWAT, *Tnfα* and *Il6* expression was substantially increased in *Bmal2*^-/-^ mice fed HFD compared to WT (Figure 3B). These results indicate that *Bmal2* invalidation amplifies the expression of specific inflammatory markers during diet-induced obesity in both liver and EpiWAT.

To further assess potential circadian regulation of these inflammatory genes, we monitored their expression over 24-hour-light-dark cycle in both ND and HFD fed mice (Figure 3C-F). In the liver, only minor changes were observed in the circadian expression profiles of *Tnfα* and *Il6* between *Bmal2*^-/-^ and WT mice under both diets (Figure 3C, D). In contrast, *Tnfα* circadian expression in EpiWAT was altered in *Bmal2*^-/-^ mice when compared to WT even under ND, evidenced by significant modifications in MESOR and phase cosinor parameters (Figure 3E). Under HFD, the *Tnfα* circadian expression was further disrupted in *Bmal2*^-/-^ mice compared to WT. Meanwhile, the *Il6* circadian expression pattern in EpiWAT was not significantly disrupted by *Bmal2* invalidation (Figure 3F).

Overall, these findings demonstrate that the absence of *Bmal2* increases inflammatory responses in both liver and EpiWAT, particularly under HFD-induced metabolic stress, likely contributing to the altered metabolic phenotype observed in obese *Bmal2*^-/-^ mice.

### 3.4. *Bmal2* invalidation promotes CD8^+^ T cell infiltration in liver and visceral adipose tissue under high fat diet

Given the exacerbated inflammatory state observed in *Bmal2*^-/-^ mice fed an HFD, we next assessed whether alterations in immune cell composition underlie this phenotype. We quantified the frequencies of macrophages and T lymphocytes in both liver and EpiWAT of *Bmal2*^-/-^ and WT mice fed either a ND or HFD. Under ND, *Bmal2*^-/-^ mice exhibited a significant increase in double-negative CD4^-^CD8^-^ (DN) T cells in the liver (Supp.Figure 3A) and in CD4^+^ T cells in EpiWAT (Supp.Figure 3B), relative to WT. However, no significant difference in macrophage frequency was observed between *Bmal2*^-/-^ and WT in either tissue (Supp.Figure 3C-G). In HFD condition, total macrophages (F4/80^+^) frequency did not differ between in *Bmal2*^-/-^ WT mice in either liver or EpiWAT (Figure 4A-B).

**Fig. 4.**
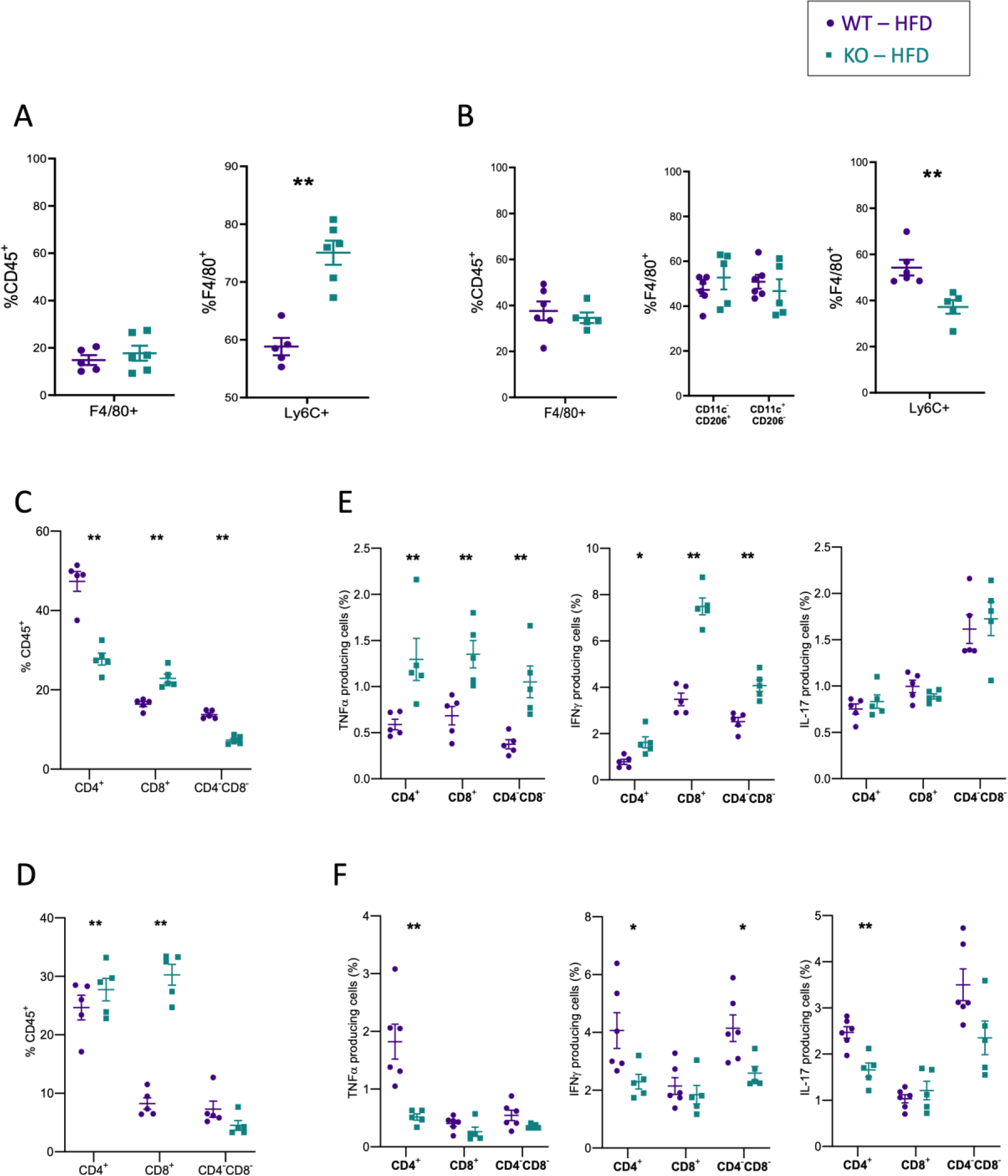
Impaired immune cell phenotypes in liver and EpiWAT *of obese Bmal2^-/-^*mice. A) Frequency of F4/80^+^ cells among CD45^+^ cells and Ly6C^+^ among F4/80^+^ cells in liver (%, n=5-6 per group), B) Frequency of F4/80^+^ cells among CD45^+^ cells, CD11c^-^ CD206^+^ and CD11c^+^CD206^-^ among F4/80^+^ cells, and Ly6C^+^ among F4/80^+^ cells, in EpiWAT (%, n=5-6 per group); Frequency of CD4^+^, CD8^+^ and CD4^-^CD8^-^ T cells among CD45^+^ cells in (C) liver and (D) EpiWAT (n=5 per group), and frequency of TNFα, IFNγ and IL-17-producing CD4^+^, CD8^+^ and CD4^-^CD8^-^ T cells in (E) liver and (F) EpiWAT (n=5 per group) from *Bmal2^-/-^*mice (KO) and controls (WT) fed with HFD during 12 weeks. All data are represented as mean ± S.E.M. All statistical analyses were performed by two-tailed Mann–Whitney test. *P < 0.05, **P < 0.01, ***P < 0.001.

Nevertheless, the proportion of F4/80^+^ Ly6C^+^ cells was increased in liver whereas it was decreased in the EpiWAT of *Bmal2*^-/-^ HFD-fed mice when compared to WT mice (Figure 4A-B). T lymphocytes profiling in the liver showed a reduction in CD4^+^ and DN T cells concomitant with an increase in CD8^+^ T cells in *Bmal2*^-/-^ mice fed HFD (Figure 4C). Meanwhile in EpiWAT, both CD4^+^ and CD8^+^ T cell frequencies were elevated in *Bmal2*^-/-^ mice relative to WT (Figure 4D).

To elucidate whether these shifts in T cell populations correlate with changes in their inflammatory potential, we measured cytokine production by conventional T lymphocyte subsets in both liver and EpiWAT. In the liver of HFD-fed *Bmal2*^-/-^ mice, TNFα- and IFNγ-producing CD4^+^, CD8^+^ and DN T cells were significantly more abundant compared to WT, whereas IL-17-producing cells did not differ (Figure 4E, Supp.Figure 4A). However, the frequencies of TNFα, IFNγ or IL-17-producing T cells (either CD4^+^ T, CD8^+^ or DN) were decreased or unchanged in EpiWAT *Bmal2*^-/-^ HFD-fed mice relative to WT (Figure 4F, Supp.Figure 4B). These results support the notion that the characteristic inflammatory environment in the liver of *Bmal2*^-/-^ mice in obesogenic condition is predominantly driven by inflammatory T lymphocytes, whereas the adipose tissue inflammation may largely originate from the non-immune fraction itself with immune cell infiltration playing a comparatively smaller role in EpiWAT.

### 3.5. Loss of *Bmal2* enhances loss of adipose progenitor identity during diet-induced obesity

To clarify whether *Bmal2* deficiency impacts adipose tissue at the level of mature adipocytes, we isolated EpiWAT adipocytes from *Bmal2*^-/-^ and WT mice under ND and HFD conditions. Transcript analysis of inflammatory genes (*Tnfα, Il6*) (Figure 5A-B) revealed that even under ND, *Tnfα* expression was significantly upregulated in *Bmal2*^-/-^ adipocytes compared to WT (Figure 5B). Under HFD, both *Tnfα* and *Il6* levels were markedly higher in *Bmal2*^-/-^ adipocytes relative to WT (Figure 5B). Notably, the transcript level of *Nfkb1*, which encodes a central transcription factor for *Tnfα*, was also elevated in *Bmal2*^-/-^ adipocytes under HFD (Supp.Figure 5A).

**Fig. 5.**
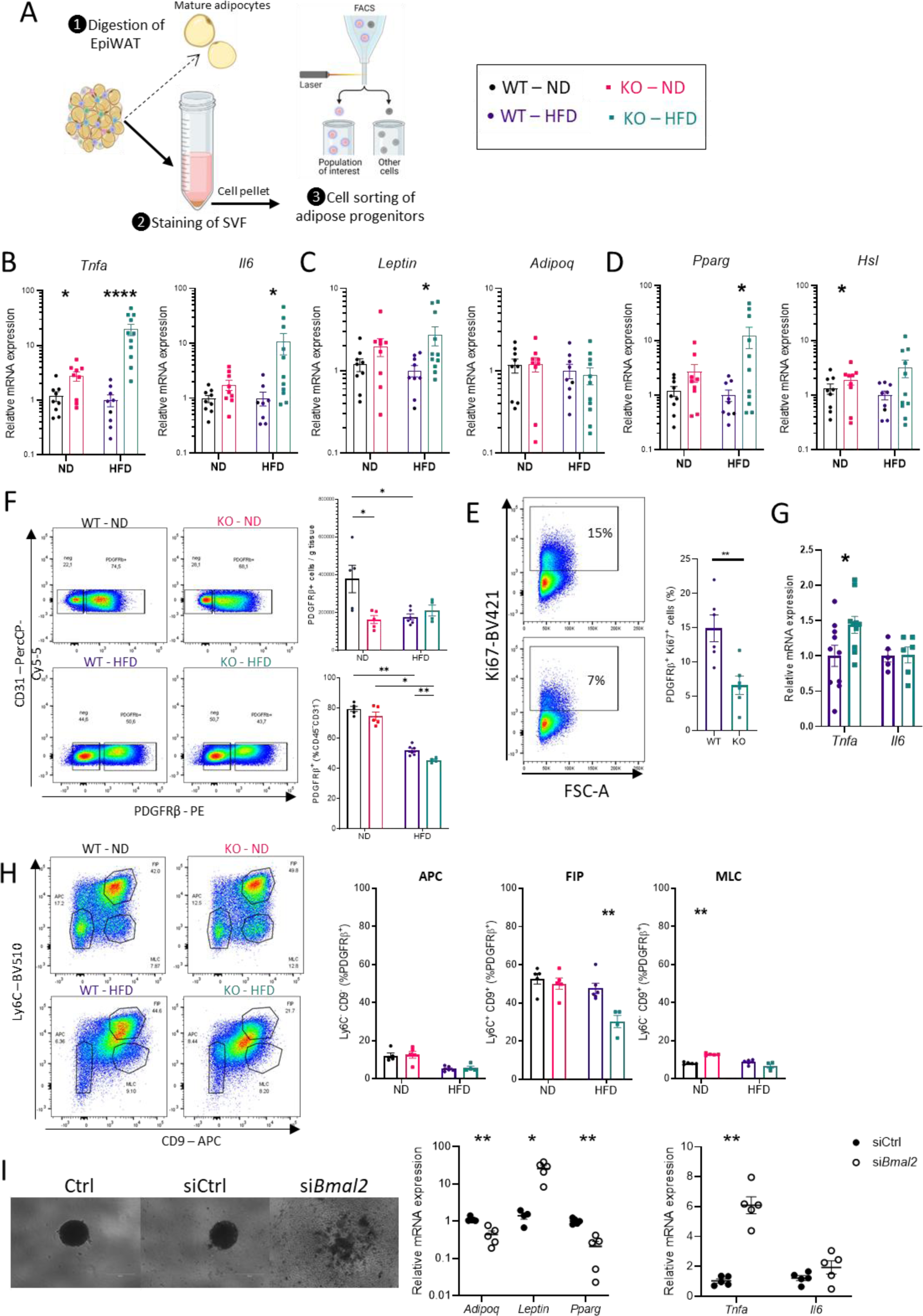
*Bmal2* invalidation enhances inflammation and impairs proliferation in PDGFRβ^+^ adipose progenitors during diet-induced obesity. A) Experimental design used to isolate mature adipocytes and adipose progenitors from murine EpiWAT. Adipocytes and PDGFRβ^+^ adipose progenitors were isolated from EpiWAT of *Bmal2^-/-^*mice (KO) and controls (WT) fed by ND or 12 weeks HFD. RT-qPCR analysis of (B) *Tnfa* and *Il6, (C) Leptin and Adipoq, and (D) Pparg and Hsl* expression in mature adipocytes (n=9-11 per group). E) Representative dot plots and quantification of PDGFRβ^+^ cells per gram of tissue (upper panel) and PDGFRβ^+^ frequency among CD31^-^CD45^-^ cells (lower panel) (n=4-6 per group). F) Representative dot plots and quantification of PDGFRβ^+^Ki67^+^ cells among PDGFRβ^+^ cells (n=5-6 per group). G) RT-qPCR analysis of *Tnfa* and *Il6* expression in PDGFRβ^+^ progenitors (n=5-10 per group). H) Representative dot plots and quantifications of PDGFRβ^+^ adipose progenitors: Ly6C^-^CD9^-^cells (APC), Ly6C^+^CD9^+^ cells (FIP) and Ly6C^-^CD9^+^cells (MLC) (n=4-6 per group). I) *in vitro* spheroid model of adipose tissue based on primary EpiWAT progenitors without treatment (Ctrl), treated with siRNA control (siCtrl) or siRNA targeting *Bmal2* (si*Bmal2)*: representative pictures (left) and RT-qPCR analysis (right) of *Adipoq*, *Leptin*, *Pparg*, *Tnfa* and *Il6* expression. Data are represented as mean ± S.E.M. Statistical analyses were performed by two-tailed Mann-Whitney test. *P < 0.05, **P < 0.01, ***P < 0.001.

Analyses of metabolic markers expression indicated that *Bmal2*^-/-^ adipocytes exhibit functional disruptions even in basal conditions: *Hsl* (hormone-senstive lipase) expression was slightly but significantly increased (Figure 5D), and both *Leptin* and *Pparg* expression tended to be higher in *Bmal2*^-/-^ adipocytes than in WT (Figure 5C-D), suggesting an enhanced lipogenic and lipolytic potential linked to chronic inflammation.

Next, we investigated whether a similar metabolic and inflammatory dysfunction could bemeasured in adipose progenitors prior to differentiation. As previously described^61^, we isolated CD45^-^CD31^-^PDGFRβ^+^ cells from the EpiWAT (see Figure 5A). Previous studies have shown that HFD generally reduces the frequency of adipose progenitors^61,62^ but increases total cell count per fat pad in C57BL/6 mice^62^. In agreement with these findings, we observed a decreased proportion of PDGFRβ^+^ progenitors per gram of tissue in HFD versus ND-fed WT mice, as well as in *Bmal2*^-/-^ ND fed mice compared to WT-ND fed mice (Figure 5E; Supp.Figure 5B). The decline in PDGFRβ^+^ progenitor frequency was even more pronounced during HFD in *Bmal2*^-/-^ mice compared to WT (Figure 5E), suggesting that *Bmal2* deficiency mirrors the detrimental effect of HFD on progenitors cells and that combining *Bmal2* invalidation with HFD exacerbates these impairments. Consistent with this, *Bmal2^-/-^* HFD-fed mice showed a significant reduction in PDGFRβ^+^ Ki67^+^ cell frequency, indicative of decreased proliferation capacity (Figure 5F).

We further examined whether these progenitor cells exhibit early inflammatory alterations. PDGFRβ^+^ progenitors from HFD fed *Bmal2*^-/-^ mice expressed higher *Tnfα* levels than those from WT (Figure 5G), indicating an inflammatory shift at a pre-adipocyte stage. No significant difference in *Tnfα* expression was observed in ND condition (Supp. Figure 5C).

To gain deeper insights into progenitor subpopulations, we assessed CD9/Ly6C markers, classifying cells into adipogenic pre-adipocytes (APC: CD9^-^Ly6C^-^), fibrogenic and inflammatory progenitors (FIP : CD9^+^Ly6C^+^), and mesothelial-like cells (MLC: CD9^+^Ly6C^-^) as previously described^61^. Under ND, WT mice displayed distinct progenitor subsets, whereas HFD-fed WT mice had more homogeneous progenitor population (Figure 5H). Remarkably, *Bmal2*^-/-^ HFD-fed mice showed even greater alteration in these progenitor subpopulations compared to WT controls, suggesting that *Bmal2* deficiency compounds the HFD-driven shift in progenitor identity.

Finally, to test the ability of *Bmal2*^-/-^ progenitors to sustain normal adipose tissue function, we employed an *in vitro* spheroid model using primary adipose tissue progenitors combined with *siBmal2* (Figure 5I). Silencing *Bmal2* markedly reduced spheroid formation compared to both the untreated control and siControl, alongside decreased expression of key adipocytes functional markers (*Adipoq, Leptin, Pparg*) and a pronounced elevation of *Tnfa* and *Il6* expression.

Overall, *Bmal2* invalidation enhances both inflammation and metabolic dysfunction in EpiWAT adipocytes and their progenitors, likely impairing the tissue capacity to adapt to diet-induced obesity. These findings strongly suggest that BMAL2 is a critical regulator of adipocytes inflammatory response and of adipose progenitor identity maintenance.

### 3.6. Invalidation of *Bmal2* activates apoptosis and inflammation transcriptional programs in adipose progenitors during diet-induced obesity

To delineate the transcriptomic changes in adipose progenitors due to *Bmal2* deficiency, we performed bulk RNA sequencing of PDGFRβ^+^ progenitors isolated form EpiWAT. Principal component analysis (PCA) of these samples revealed a clear two-dimensional separation between *Bmal2*^-/-^ and WT groups in both ND and HFD conditions (Figure 6A). Despite this consistent segregation, the genes driving the divergence differed notably between ND and HFD cohorts (Figure 6B), indicating distinct molecular signatures shaped by diet.

**Fig. 6.**
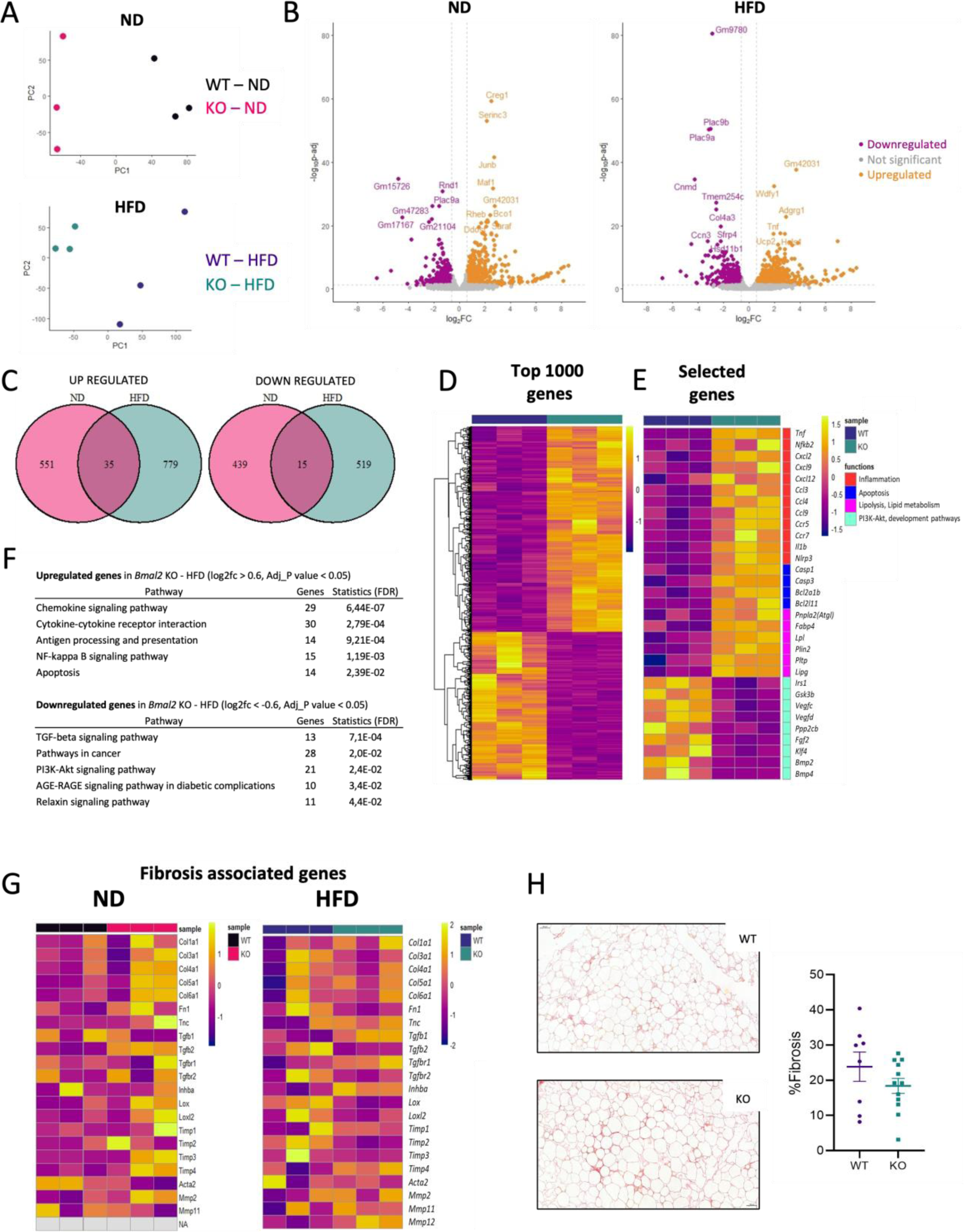
*Bmal2* invalidation induces changes of the transcriptome of PDGFRβ^+^ adipose progenitors during diet-induced obesity towards inflammatory and apoptotic gene signalling pathways. A) PCA plots showing distance between PDGFRβ^+^ adipose progenitors between WT and *Bmal2^-/-^* mice (KO) fed by ND (upper panel) or 12 weeks HFD (lower panel). B) Volcano plots highlighting significant differentially expressed genes in PDGFRβ^+^ adipose progenitors of WT and *Bmal2^-/-^* mice (KO) fed by ND (left) or 12 weeks HFD (right). C) Venn-diagram showing common up and down regulated genes PDGFRβ^+^ adipose progenitors of WT and *Bmal2^-/-^* mice (KO) fed by ND or 12 weeks HFD. D) Heatmaps showing the top 1000 differentially expressed genes and (E) selection of 31 differentially expressed genes in PDGFRβ^+^ adipose progenitors of WT and *Bmal2^-/-^* mice (KO) fed 12 weeks with HFD. F) Selection of DAVID pathways analysis of significantly upregulated and downregulated genes (P<0,05), G) selected fibrosis associated genes expressed in PDGFRβ^+^ adipose progenitors of WT and *Bmal2^-/-^* mice (KO) fed 12 weeks with HFD. Differentially expressed genes were defined with log2FC>0.6 and p-adj<0.05. n=3 per group. H) Histology analysis of the EpiWAT of HFD fed mice using Sirius red staining and quantification (n=8-11). Data are represented as mean ± S.E.M. All statistical analyses were performed by two-tailed Mann-Whitney test. *P < 0.05, **P < 0.01, ***P < 0.001.

In ND condition, 1040 genes were differentially expressed (log2FC>0.6 and p-adj<0.05) between *Bmal2*^-/-^ and WT adipose progenitors (587 upregulated, 454 downregulated) (Figure 6B). In HFD condition, a total of 1348 genes were differentially expressed (814 upregulated, 534 downregulated) (Figure 6B). Notably, *Tnfα* ranked among the most upregulated genes in PDGFRβ^+^ progenitors from *Bmal2*^-/-^ HFD-fed mice. The larger number of differentially expressed genes in HFD-fed mice underscores the interaction between *Bmal2* invalidation and obesogenic stress.

A comparative analysis of genes differentially expressed in *Bmal2*^-/-^ progenitors under ND and HFD reveled limited overlap. Specifically, 35 genes were commonly upregulated irrespective of diet, 551 were uniquely upregulated under ND, and 779 genes were uniquely upregulated in HFD (Figure 6C). Likewise, 15 genes were commonly downregulated irrespective of diet, 439 genes were uniquely downregulated under ND and 519 genes were uniquely downregulated under HFD (Figure 6C). This pattern indicates that *Bmal2*^-/-^ adipose progenitors develop distinct transcriptional responses depending on dietary conditions.

Comparison of the top 1000 differentially expressed genes showed markedly different transcriptional signature between *Bmal2*^-/-^ versus WT progenitors in HFD condition (Figure 6D). Gene ontology analysis indicated an up-regulation of inflammatory mediators, apoptotic pathways, lipolysis, and lipid metabolism in *Bmal2*^-/-^ progenitors (Figure 6E, F). Conversely, genes involved in PI3K-Akt signaling axis, cellular development and maintenance were downregulated in *Bmal2*^-/-^ mice under HFD (Figure 6E, F). Those pathways were not differentially regulated in ND condition (Supp. Figure 6A-C). Gene set enrichment analysis (GSEA, FDR 25%) validated 19 enriched upregulated pathways in the *Bmal2*^-/-^ mice, including inflammatory responses (e.g. TNFα signaling via NF_k_B, IL6-JAK-STAT3 signaling) and apoptosis, as well as one downregulated metabolic pathway (oxidative phosphorylation) under HFD (Supp. Figure 6D).

Since increased inflammation can drive fibrosis and that adipose progenitors have been found to shift to a profibrogenic phenotype during obesity^62^, we investigated fibrosis markers in *Bmal2*^-/-^ EpiWAT progenitors. Neither RNA-seq data (Figure 6G) nor histological analyses of EpiWAT (Figure 6H) showed significant differences in expression of profibrogenic genes by PDGFRβ^+^ progenitors or tissue fibrosis between *Bmal*2^-/-^ and WT mice, under either ND or HFD.

Overall, these findings demonstrate that *Bmal2* invalidation strongly reshapes the transcriptome of adipose progenitors and, in conjunction with an obesogenic diet, amplifies deleterious effects on progenitor function. Specifically, loss of *Bmal2* enhanced inflammatory and apoptotic transcriptional programs while compromising metabolic and maintenance programs in adipose progenitors, which might limit their adaptive capacity during diet-induced obesity.

## 4. DISCUSSION

Chronic disruption of circadian rhythms is increasingly recognized as a key contributor to obesity and its associated comorbidities, highlighting the need for a deeper understanding of how these biological clocks regulate energy homeostasis, inflammation and tissue remodeling. In the present study, we investigated the role of the circadian clock gene *Bmal2* in modulating the immune-metabolic adaptation of adipose tissue during obesity. Our findings reveal that *Bmal2* regulates inflammatory processes in EpiWAT through adipocytes, adipose progenitors, and immune cells, exerting systemic effects that also compromise liver metabolism and insulin sensitivity.

In the obesity-prone C57BL/6 mouse model, we observed an increased *Bmal2* expression in the EpiWAT of WT mice during HFD feeding. By contrast, *Bmal2*^-/-^ mice displayed exacerbated weight gain under the same obesogenic condition, accompanied by extensive immune-metabolic dysfunction in EpiWAT, particularly marked by elevated *Tnfα* expression, as well as hepatic steatosis and insulin resistance. These data suggest that *Bmal2* acts as a crucial regulatory node, linking adipose tissue inflammation to liver metabolism. In addition to the previously reported changes in circadian regulation of feeding and energy balance in *Bmal2^-/-^* mice^60^, the significant disruption of the *Tnfα* circadian expression pattern in the EpiWAT of *Bmal2^-/-^* lean and obese mice suggests that disturbances in the circadian system at both the adipose tissue and central nervous system levels may cause or at least contribute to their altered phenotype.

Interestingly, the phenotype of *Bmal2*^-/-^ mice under ND was comparatively mild, notably in terms of tissue-specific inflammatory gene expression. This findings echoes our earlier work on the role of *Bmal2* in transcriptional regulation of *Il21* in type 1 diabetes context, in which *Il21* induction by BMAL2 was highly dependent on inflammatory and developmental cues in younger animals^59^. The modest phenotype we observed in ND-fed *Bmal2*^-/-^ mice implies that BMAL2-dependent transcription way require specific chromatin or metabolic configurations to activate inflammatory pathways^63^. In line with this concept, we detected no prominent difference in *Il21* expression under HFD in the current study, suggesting that the influence of BMAL2 on *Il21* expression may be age or context dependent.

Our data further indicate that BMAL2 helps to maintain EpiWAT homeostasis by preventing excessive inflammation and safeguarding the progenitor niche. The pronounced upregulation of *Tnfα* in *Bmal2*^-/-^ adipocytes and progenitors points to the potential for direct or indirect transcriptional repression of *Tnfα* by BMAL2. Notably, previous *in vitro* studies have shown that BMAL2 expression is induced by TNF-α^64,65^, supporting the hypothesis of a reciprocal relationship in which BMAL2 and TNF-α form a regulatory loop controlling adipose tissue inflammation. Consequently, the insulin resistance observed in *Bmal2*^-/-^ mice under HFD conditions may stem predominately from enhanced inflammatory signaling, mediated by TNF-α, IL-6 and related cytokines, which disrupt the insulin signaling cascade.

The limited alterations in immune cells phenotype within the EpiWAT of *Bmal2*^-/-^ mice shifted our attention toward adipocytes and adipose progenitors. We found that *Bmal2*^-/-^ EpiWAT displayed reduced lipid storage, concurrent with hypertrophic adipocytes and impaired adipogenesis. Mature adipocytes and PDGFRβ^+^ progenitors from *Bmal2*^-/-^ mice showed elevated expression of proinflammatory cytokines, especially TNF-α, aligning with previous reports linking circadian disruption in adipocytes and progenitors to increased inflammation during obesity^9^. Other studies have reported an expansion of fibrogenic and inflammatory progenitors (FIP) and a decreased adipogenic progenitors (APC) in diet-induced obesity^61,62^. In contrast, *Bmal2* deficiency reshaped adipose progenitor populations differently, diminishing FIP cells while modifying APC population toward a more uniform state with reduced proliferative and metabolic capacities. This remodeling potentially denotes a loss of ability to diversify into distinct functional lineages, a phenomenon reminiscent of disrupted circadian regulation of adipose progenitor proliferation^27^. Thus, in an obesogenic environment, defective BMAL2 activity may bias progenitors toward apoptosis and inflammatory phenotype rather than promoting adaptive tissue expansion and metabolic response.

An emerging body of work underscores circadian regulation of adipogenesis via multiple pathways^30,32,54,66–68^. For instance, *Bmal2* expression reportedly increase during early adipocyte differentiation *in vitro*, yet can also repress adipogenesis by targeting transcription factors such as BMAL1, KLF15 and C/EBPβ^54^. Consistent with these findings, we detected increased *Bmal2* expression in EpiWAT of HFD-fed WT mice, pointing to its involvement in adipogenesis under obesogenic conditions. Further investigations, including *in vitro* differentiation of preadipocyte cell line (3T3-L1, 3T3-F442A) with *Bmal2* knockdown or overexpression could help delineate the mechanisms through which BMAL2 integrates circadian control with adipocyte differentiation.

Obesity induces a state of chronic stress in adipocytes, shifting them toward a pro-inflammatory profile and impairing their capacity for lipogenesis^69^. In line with this, *Bmal2*^-/-^ adipocytes exhibited heightened inflammation and diminished lipid storage. Understanding how BMAL2 modulates immune and metabolic gene networks in discrete adipocyte subpopulations will provide a more granular view of the role of circadian regulation in adipose tissue.

Notably BMAL1 also exerts significant control over adipogenesis and lipid handling. However, whether BMAL1 primarily activates or represses adipocyte differentiation remains debated. Although BMAL1 regulates inflammatory mechanisms in immune cells and adipose tissue^9,41,44,70^ and that *Bmal2*^-/-^ and *Bmal1^-/-^* mice share certain phenotypic features, they also exhibit distinct alterations in weight gain, lipid storage distribution, and insulin secretion capacity^30,31,34^. The present findings emphasize the necessity of discussing the individual contributions of these clock genes in orchestrating adipose tissue plasticity via progenitors and mature adipocytes, especially in the context of obesity.

Finally, while *BMAL2* has been linked to metabolic disease in human genetic association studies ^52,53^, our mouse findings strongly suggest that *Bmal2* deletion disrupts adipose-liver crosstalk, leading to ectopic lipid deposition in the liver and the onset of insulin resistance. Tissue specific *Bmal2* deletion in adipocytes or hepatocytes could provide critical insights into how this circadian regulator modulates systemic metabolism by coordinating the interplay between adipose and hepatic tissues.

In conclusion, our study demonstrates that BMAL2 is an essential regulator of immune-metabolic adaptation in adipose tissue, particularly in obesogenic conditions. By modulating inflammatory gene expression and progenitor function, *Bmal2* deletion exacerbates adipose tissue dysfunction, reduces EpiWAT lipid storage capacity, and promotes insulin resistance in both adipose tissue and liver. These findings underscore the intricate relationship between circadian regulation, adipose tissue plasticity, and metabolic homeostasis, thereby opening new avenue for therapeutic strategies targeting circadian pathways to combat obesity and its complications.

## List of Supplementary Materials

Fig S1 to S6

Tables S1 to S2

## Acknowledgments

We thank Cybio, HistIM and Genom’IC facilities of the Institut Cochin. We thank all the staff of Pasteur Animal Facility, and in particular Myriam Mattei for taking care of the animals during the COVID-19 lockdown. We thank Christian Boitard for support, and Sandrine Luce, Latif Rachdi and Masaya Oshima for their scientific and technical advice.

## Fundings

This work was supported by current fundings from Inserm, CNRS, and Université Paris Cité, individual grants from Association Française pour l’Etude du Foie (AFEF 2023), Société Francophone du Diabète (SFD 2023) to A.T., European Foundation for the Study of Diabetes to A.L. and A.T, Laboratoire d’Excellence consortium Inflamex (ANR-11-IDEX-0005-02), Recherche Hospitalo-Universitaire (RHU) QUID-NASH program (ANR-17-RHUS-009), Fondation pour la Recherche Médicale (FRM EQU201903007779), Agence National de Recherche (ANR-19-CE14-0041-01 HEPADIMAIT) to A.L., DHU authors to U.R., M.P. and B.F. were supported by the French Ministry of Research.

## Author contributions

M.P. and A.T. performed most of the experiments and data analysis; B.F., L.C., A.G., L.B. and U.R. performed experiments. M.P. performed bioinformatic analysis. M.P., A.L., U.R. and A.T. wrote the paper. N.V. and E.C. edited the paper and provided biological material. A.L., U.R. and A.T. provided scientific guidance. U.R. and A.T. supervised the work.

## Conflict of interest

The authors have declared that no conflict of interest exists.

## Data and material availability

Bulk RNA-seq data will be made available in GEO database.

## Notes

### Competing Interest Statement

The authors have declared no competing interest.

