## Supplementary figures and tables for "BMAL2 controls adipose tissue inflammation and metabolic adaptation during obesity"

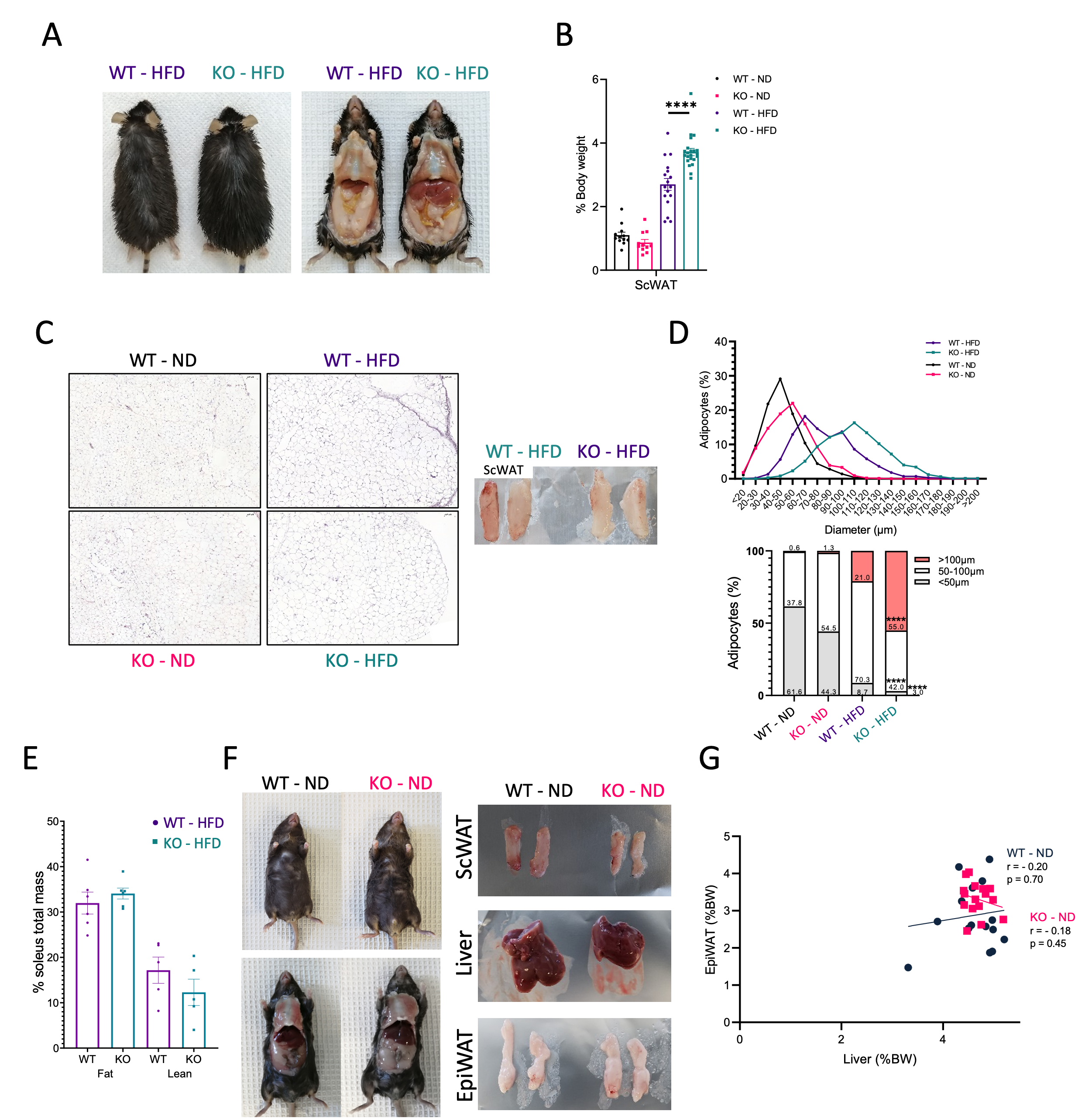

**Supp.Figure 1: Phenotype of normal and high fat diet fed *Bmal2^-/-^* mice.** A) Representative pictures of *Bmal2^-/-^* mice (KO) and controls (WT) fed with HFD during 12 weeks, B) Weight of ScWAT (KO-ND n=11, WT-ND n=12, KO-HFD n=22, WT-HFD n=17) relative to body weight (%), C) Representative hematoxylin & eosin staining of EpiWAT (scale bars, 200 μm) and representative pictures of ScWAT, D) Quantification of adipocyte size in ScWAT (100-400 adipocytes per section, 1 section per mouse, 4-18 mice per group), E) NMR-evaluated soleus muscle fat and lean mass (n=5-6 per group), F) Representative pictures of *Bmal2^-/-^* mice (KO) and controls (WT) fed with ND with corresponding pictures of tissues: ScWAT, liver, EpiWAT, and G) Spearman correlation between weight of EpiWAT and liver relative to body weight (%), from *Bmal2^-/-^* mice (KO) and controls (WT) fed with ND and/or HFD during 12 weeks. Data are represented as mean ± S.E.M. All statistical analyses were performed by two-tailed Mann-Whitney test. *P < 0.05, **P < 0.01, ***P < 0.001.

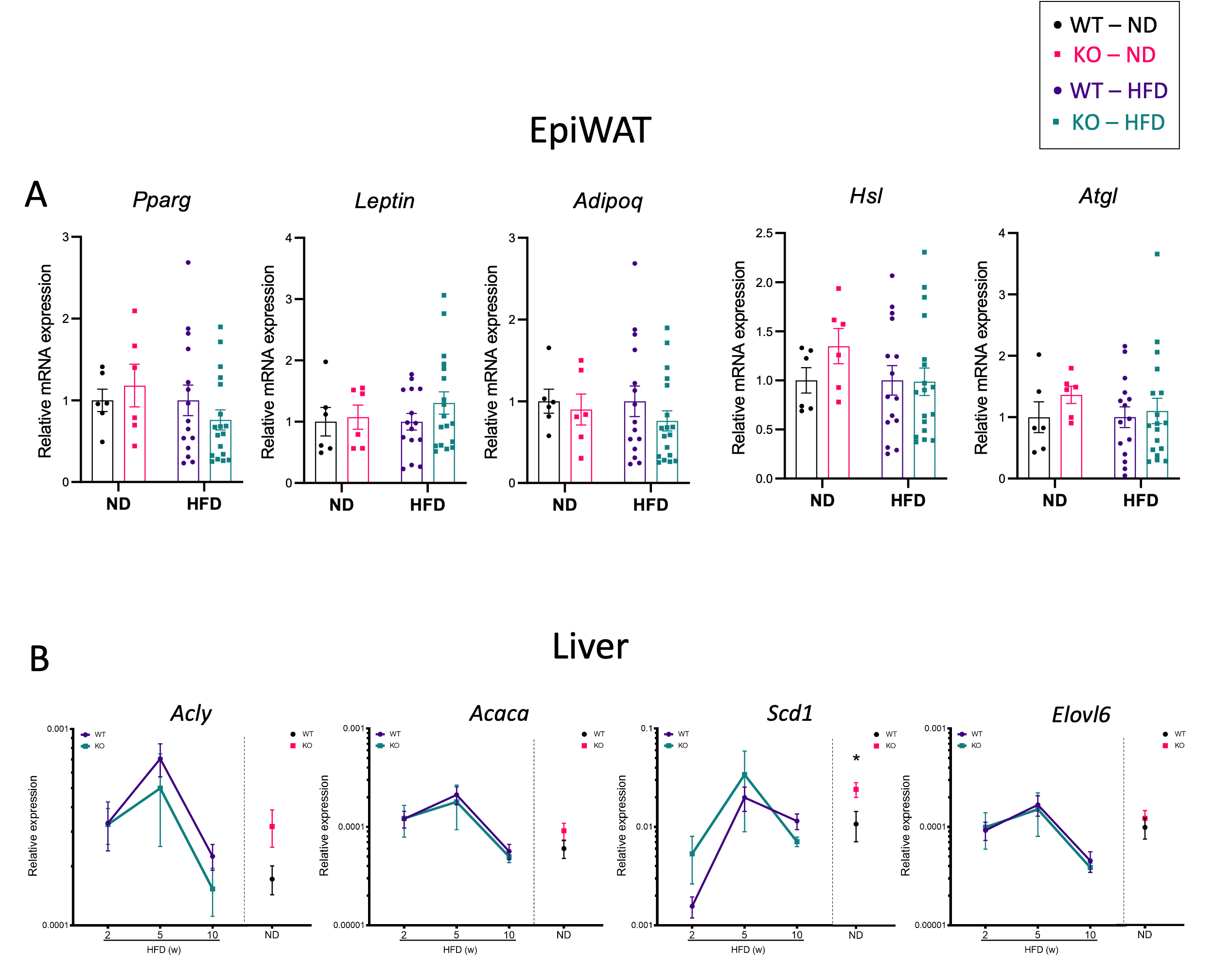

**Supp.Figure 2: Metabolic gene expressions in EpiWAT and liver from normal and high fat diet fed *Bmal2^-/-^* mice.** A) RT-qPCR analysis of *Pparg, Leptin, Adipoq, Hsl and Atgl* expression in EpiWAT (n=6-18 per group) from *Bmal2^-/-^* mice (KO) and controls (WT) fed with ND or HFD during 12 weeks. B) Kinetics of lipogenesis related genes (*Acly, Acaca, Scd1, Elovl6*) in liver from *Bmal2^-/-^* mice (KO) and controls (WT) during 2, 5 and 10 weeks of HFD and ND (equivalent age to 10 weeks HFD) (n=3-7 per group). Data are represented as mean ± S.E.M. All statistical analyses were performed by two-tailed Mann-Whitney test. *P < 0.05, **P < 0.01, ***P < 0.001.

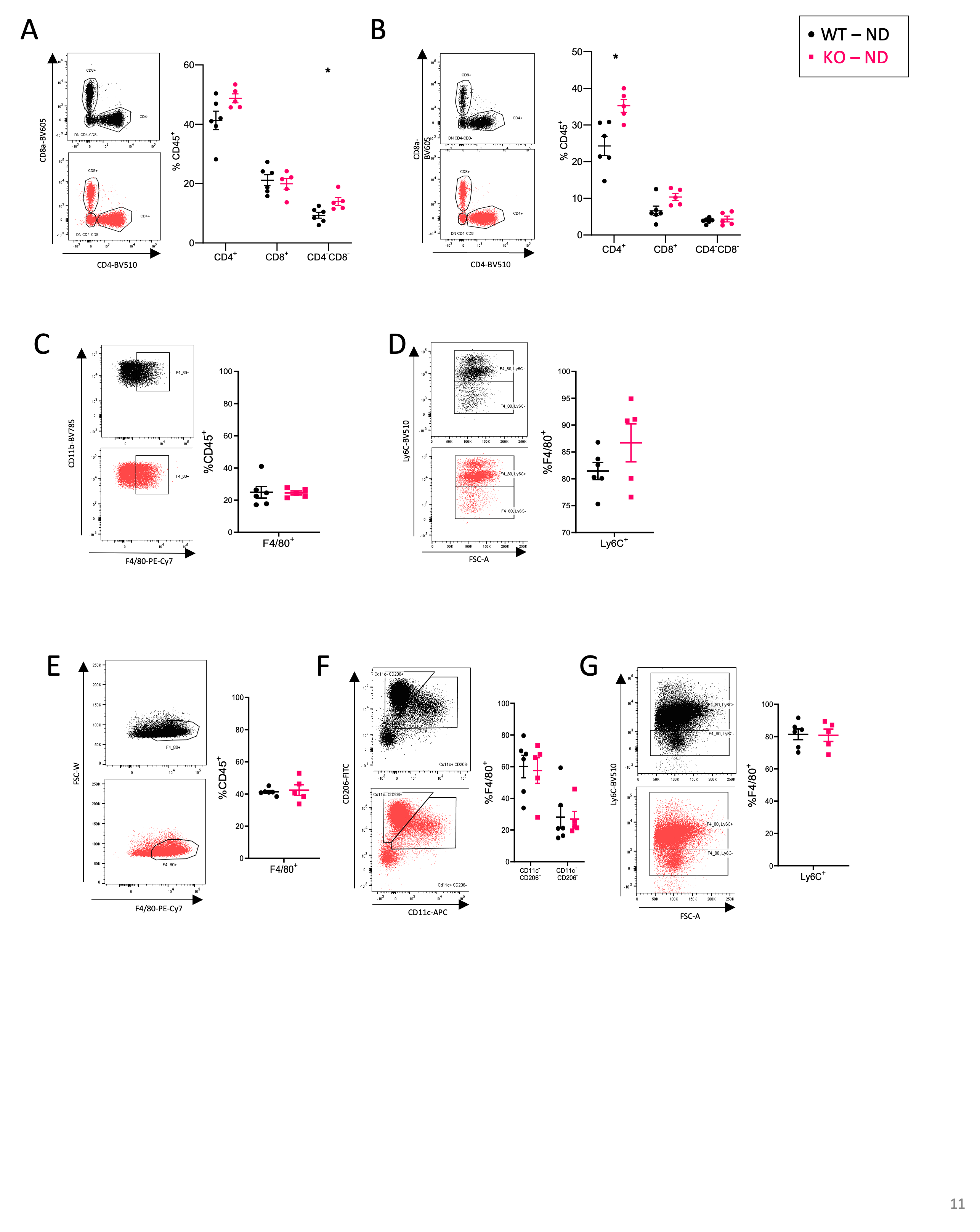
**Supp.Figure 3: Immune cell phenotype in EpiWAT and liver from normal diet fed *Bmal2^-/-^*mice**. A) Representative dot plots and frequency of F4/80^+^ cells among CD45^+^ cells, B) Ly6C^+^ among F4/80^+^ cells and C) CD4^+^, CD8^+^ and CD4^-^CD8^-^ T cells among CD45^+^ cells in liver from *Bmal2^-/-^* mice (KO, n=5) and controls (WT, n=6) fed with ND. Representative dot plots and frequency of (D) F4/80^+^ cells among CD45^+^ cells, (E) CD11c^-^CD206^+^ and CD11c^+^CD206^-^ among F4/80^+^ cells, (F) Ly6C^+^ among F4/80^+^ cells, and (G) CD4^+^, CD8^+^ and CD4^-^CD8^-^ T cells among CD45^+^ cells in EpiWAT from *Bmal2^-/-^* mice (KO, n=5) and controls (WT, n=6) fed with ND. Data are represented as mean ± S.E.M. Statistical analyses were performed by two-tailed Mann-Whitney test. *P < 0.05, **P < 0.01, ***P < 0.001.

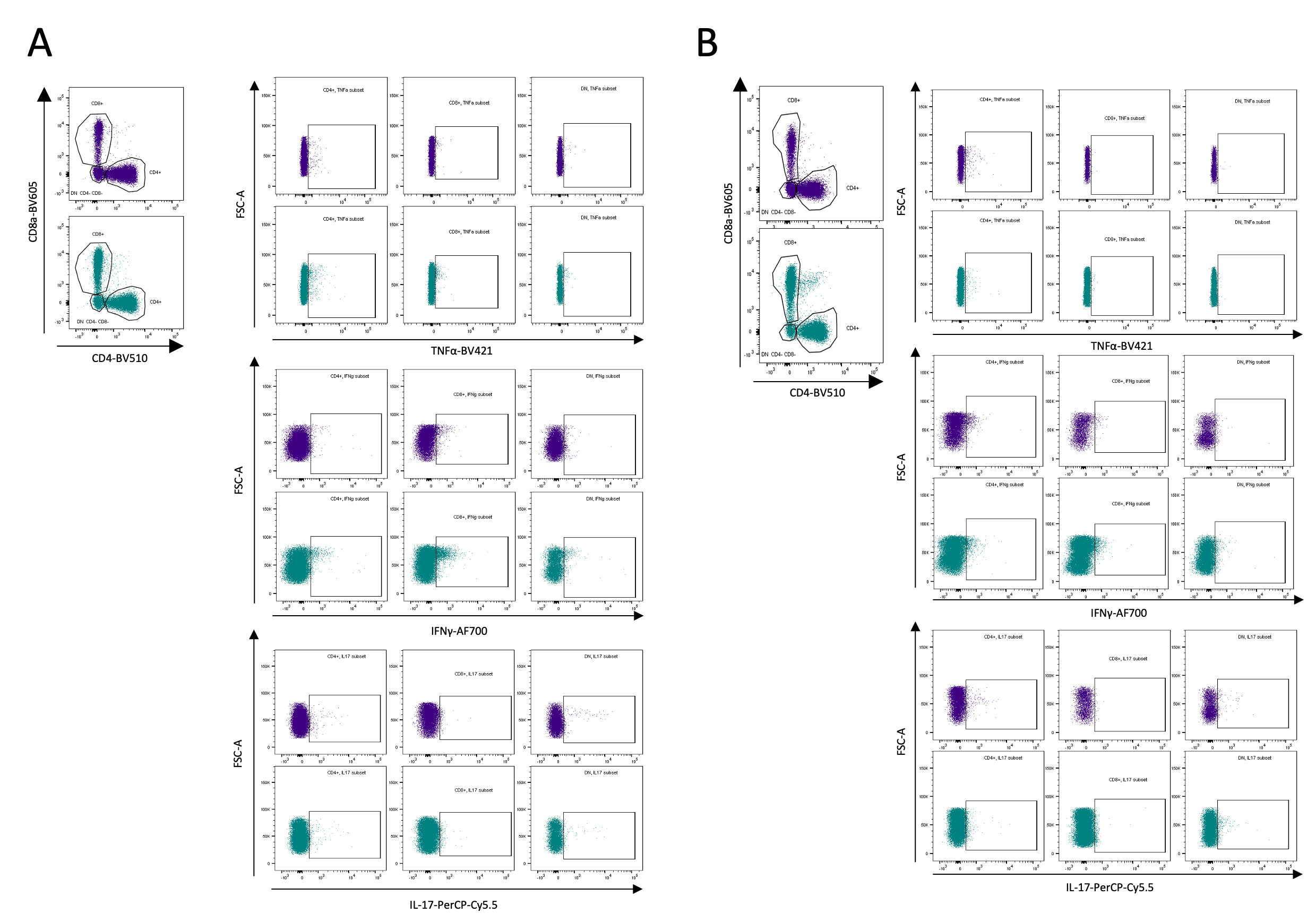
**Supp.Figure 4: Gating strategy of immune cells in EpiWAT and liver from HFD-fed *Bmal2^-/-^* mice**. Representative dot plots showing gating of A) F4/80^+^ cells among CD45^+^ cells and Ly6C^+^ among F4/80^+^ cells in liver, B) F4/80^+^ cells among CD45^+^ cells, CD11c^-^CD206^+^ and CD11c^+^CD206^-^ among F4/80^+^ cells, and Ly6C^+^ among F4/80^+^ cells, in EpiWAT, and gating of CD4^+^, CD8^+^ and CD4^-^CD8^-^ T cells among CD45^+^ cells as well as TNFα, IFNγ and IL-17-producing CD4^+^, CD8^+^ and CD4^-^CD8^-^ T cells in (C) liver and (D) EpiWAT from *Bmal2^-/-^* mice (KO) and controls (WT) fed with HFD during 12 weeks.

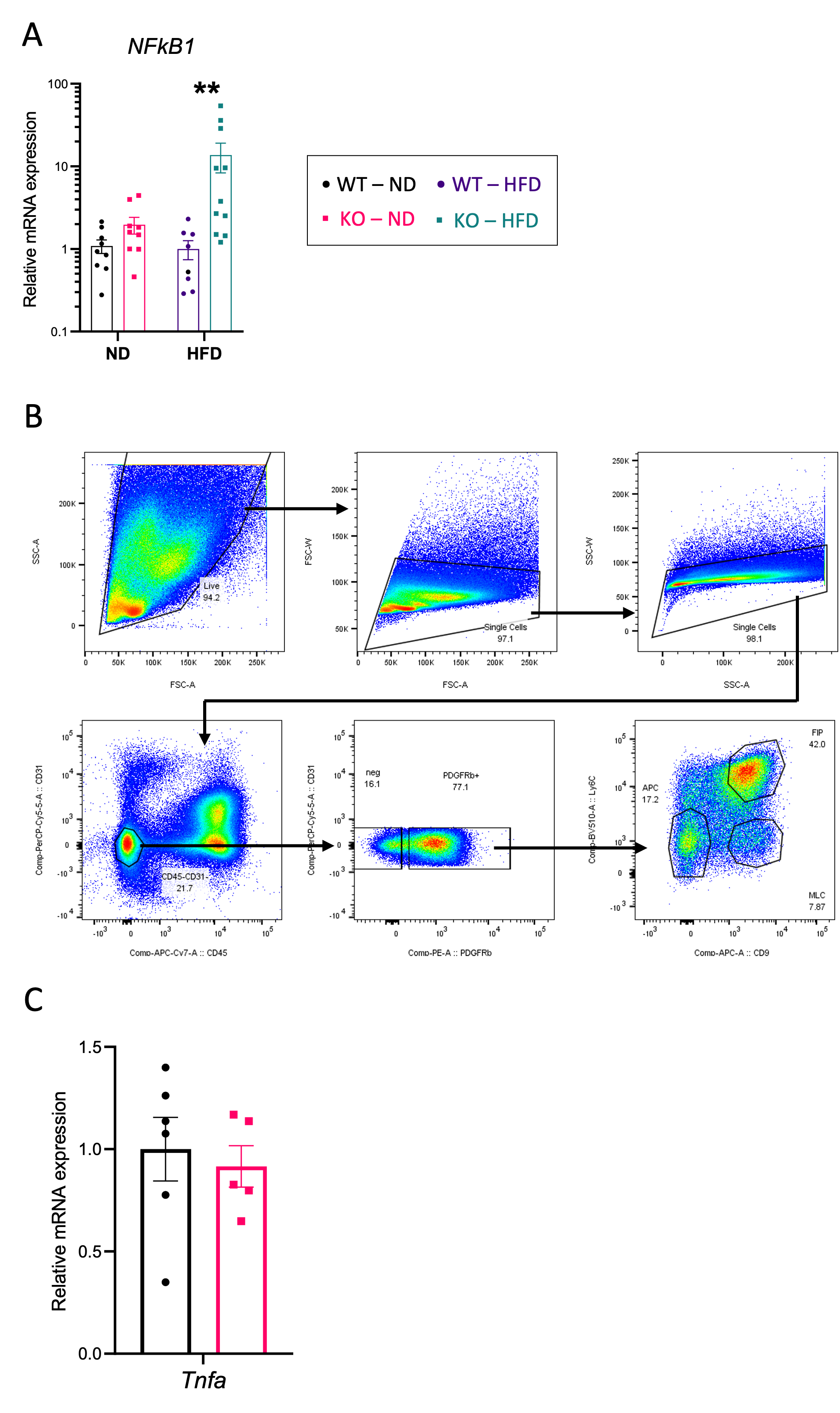
**Supp.Figure 5: Expression of *NF_k_B1* in adipocytes and gating strategy used to identify adipose progenitors in EpiWAT.** A) RT-qPCR analysis of *NF_k_B1* expression in mature adipocytes (n=8-11 per group), B) Gating strategy used to identify PDGFRβ^+^ progenitors and subpopulations: Ly6C^-^CD9^-^cells (APC), Ly6C^+^CD9^+^ cells (FIP) and Ly6C^-^CD9^+^cells (MLC), C) RT-qPCR analysis of *Tnfα* expression in PDGFRβ^+^ progenitors of ND fed mice. Adipocytes and PDGFRβ^+^ adipose progenitors were isolated from EpiWAT of *Bmal2^-/-^* mice (KO) and controls (WT) fed with ND or 12 weeks of HFD. Data are represented as means ± S.E.M. Statistical analyses were performed by two-tailed Mann-Whitney test. *P < 0.05, **P < 0.01, ***P < 0.001.

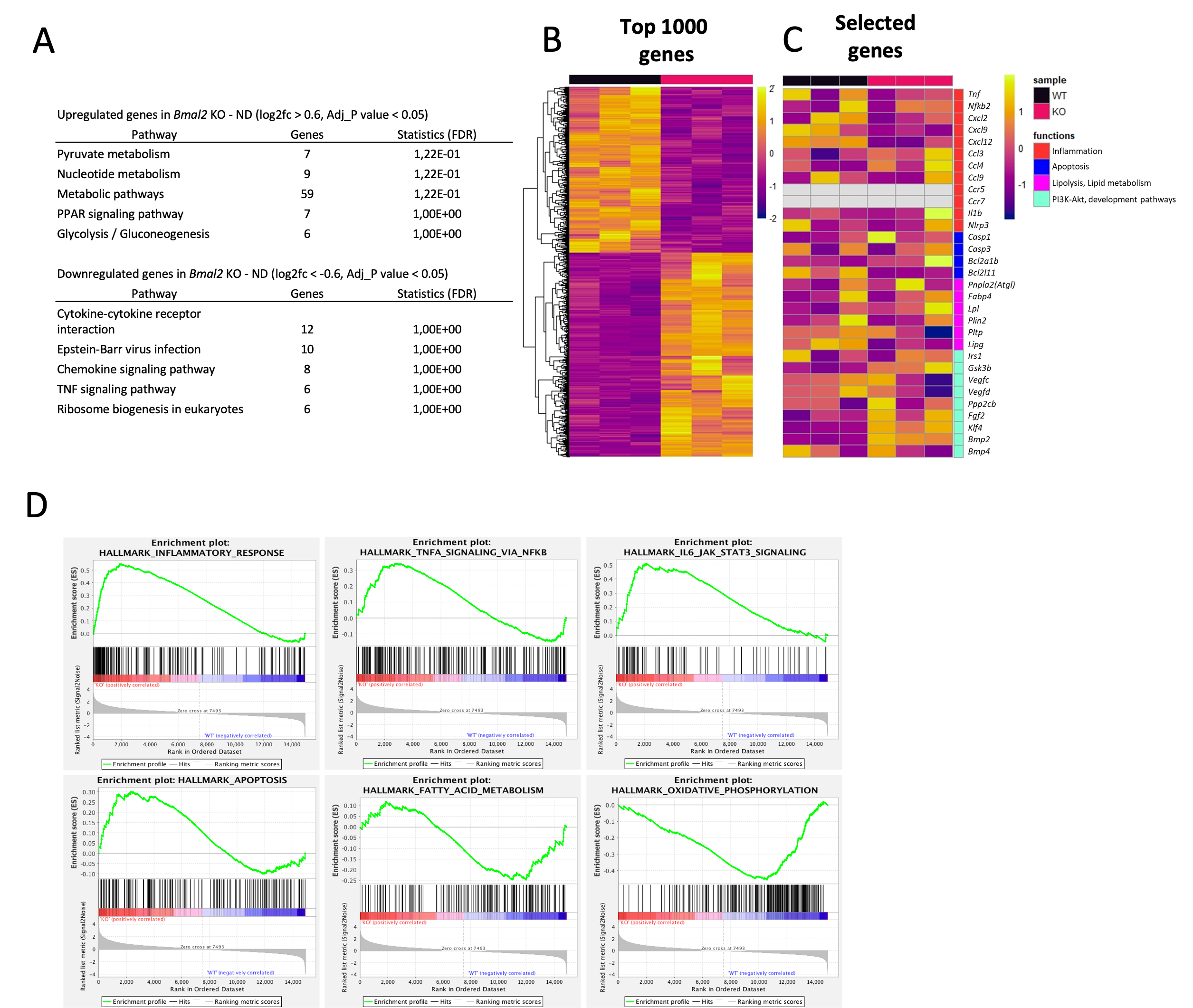

**Supp.Figure 6: RNAseq analysis under ND and gene set enrichment analysis.** A) DAVID pathways analysis of significantly upregulated and downregulated genes under ND in PDGFRβ^+^ adipose progenitors of WT and *Bmal2^-/-^* mice (KO), B) Heatmaps showing the top 1000 differentially expressed genes, and (C) selection of 31 differentially expressed genes. Differentially expressed genes were defined with log2FC>0.6 and p-adj<0.5, n=3 per group. D) Examples of the GSEA analysis of all expressed genes in PDGFRβ^+^ adipose progenitors of EpiWAT from *Bmal2^-/-^* and WT mice (murine hallmark genes, FDR 25%).

**Supplementary tables**

**Supp.Table 1: List of antibodies**

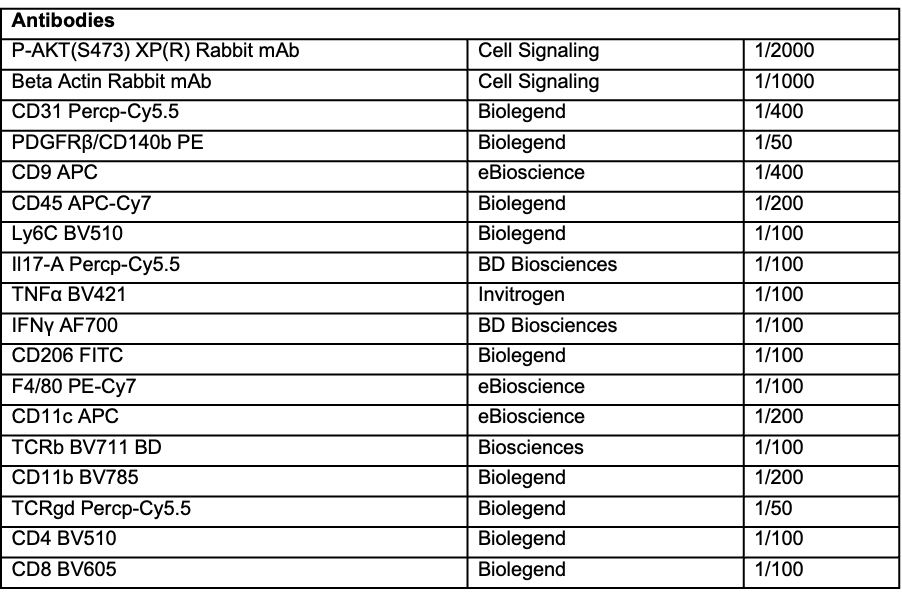

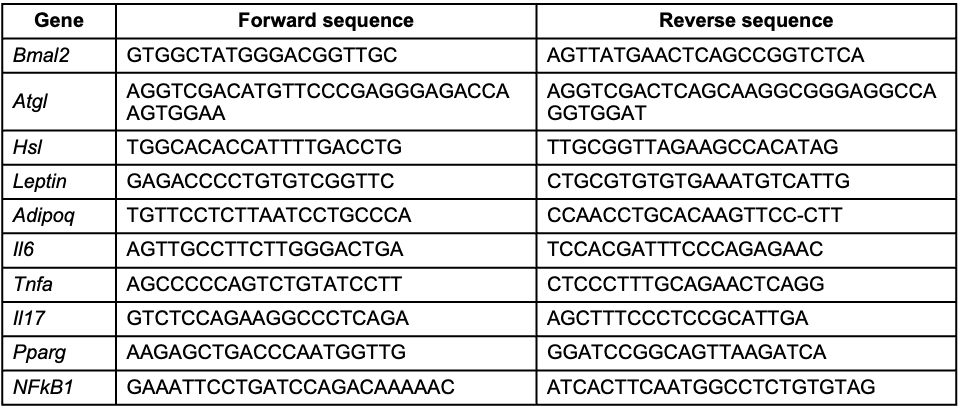
**Supp.Table 2: List of primers**
